# The cellular architecture of memory modules in *Drosophila* supports stochastic input integration

**DOI:** 10.1101/2020.12.06.413500

**Authors:** Omar A. Hafez, Benjamin Escribano, Rouven L. Ziegler, Jan J. Hirtz, Ernst Niebur, Jan Pielage

## Abstract

The ability to associate neutral stimuli with valence information and to store these associations as memories forms the basis for decision making. To determine the underlying computational principles, we build a realistic computational model of a central decision module within the *Drosophila* mushroom body (MB), the fly’s center for learning and memory. Our model combines the electron microscopy-based architecture of one MB output neuron (MBON-α3), the synaptic connectivity of its 948 presynaptic Kenyon cells (KCs), and its membrane properties obtained from patch-clamp recordings. We show that this neuron is electrotonically compact and that synaptic input corresponding to simulated odor input robustly drives its spiking behavior. Therefore, sparse innervation by KCs can efficiently control and modulate MBON activity in response to learning with minimal requirements on the specificity of synaptic localization. This architecture allows efficient storage of large numbers of memories using the flexible stochastic connectivity of the circuit.

## 1 Introduction

Operating successfully within a complex environment requires organisms to discriminate between a vast number of rewarding and stressful interactions. Learning to avoid potentially harmful interactions will promote survival of individual animals and requires long-term memory mechanisms. In recent years, significant progress has been made toward our understanding of the cellular and circuit mechanisms underlying learning and memory. Despite these advances, at the computational level it remains largely unknown how the architectures contribute to the efficient formation and storage of multiple memories.

Here, we use the *Drosophila melanogaster* mushroom body (MB) as a simple model system to investigate the computational principles underlying learning and memory in the context of decision making. A key advantage of the MB, the learning and memory center of the fly, is its relative simplicity in terms of the number and types of neurons compared to the mammalian brain. It has been demonstrated that the MB is required for the acquisition and recall of associative olfactory memories (de Belle and Heisenberg, 1994), and a wide variety of genetic tools are available to access and manipulate key cell types within the memory circuitry (Owald et al, 2015b; Lai and Lee, 2006)

Odor information is transmitted from the antennal lobe via projection neurons (PNs) to the Kenyon cells (KCs) of the MB. PNs are connected in a largely stochastic manner to the approximately 2000 KCs per brain hemisphere, resulting in a unique KC odor representation in individual flies (Litwin-Kumar et al, 2017; Aso et al, 2014a; Caron et al, 2013; Gruntman and Turner, 2013; Lin et al, 2007; Betkiewicz et al, 2020). The exception to this is a partial non-random connectivity of a population of food-responsive PNs (Zheng et al, 2022). The formation of olfactory memories within the MB circuit is assayed through experimental manipulations in which normally neutral odors are associated with either positive (reward learning) or negative (avoidance learning) valences (Livingstone et al, 1984).

Electrophysiological and calcium imaging studies demonstrated that any given odor activates approximately 3-9% of KCs (Campbell et al, 2013; Honegger et al, 2011; Turner et al, 2008; Siegenthaler et al, 2019). KCs then transmit this odor information to mushroom body output neurons (MBONs) that subdivide the MB lobes into distinct structural and functional modules by virtue of their dendritic arborizations (Owald et al, 2015a; Aso et al, 2014b; Hige et al, 2015a; Cognigni et al, 2018). During associative olfactory conditioning, the behavioral response to any given odor can be altered to encode positive (approach) or negative (avoidance) behavior. The valence of the conditioned response is encoded by the activation of dopaminergic neurons (DANs) that selectively innervate the structural MB modules defined by the MBON dendrites (Aso et al, 2014a).

DANs of the PPL1 cluster preferentially provide input to the vertical lobe of the MB and are activated during negative reinforcement, while DANs of the PAM cluster are activated during positive reinforcement and modulate activity in the horizontal lobe of the MB (Handler et al, 2019; Aso and Rubin, 2016; Tomchik and 4 Davis, 2009; Séjourné et al, 2011; Cohn et al, 2015; Burke et al, 2012). The majority of MBONs of the vertical lobe utilize acetylcholine as a neurotransmitter, and artificial activation of these MBONs often mediates approach behavior (Aso et al, 2014b). In contrast, the majority of MBONs of the horizontal lobe release glutamate as a neurotransmitter, and these neurons mediate avoidance behavior (Aso et al, 2014b).

*In vivo* calcium imaging and electrophysiological experiments demonstrated that the activity of individual MBONs can be altered by conditioning paradigms (Hige et al, 2015a; Aso and Rubin, 2016; Plaçais et al, 2013; Owald et al, 2015a; Jacob and Waddell, 2020; Perisse et al, 2016; Felsenberg et al, 2018). This modulation of MBON activity depends on the pairing of DAN and KC activity and results in either an enhancement or depression of the strength of the KC>MBON connection (Handler et al, 2019; Owald et al, 2015a; Cohn et al, 2015; Burke et al, 2012). These changes in KC>MBON connection strength occur in either the horizontal or the vertical lobe depending on the administered stimulus-reward contingencies. As a consequence, the resulting net activity of all MBONs is shifted to either approach or avoidance behavior and the behavioral response towards the conditioned stimulus during memory recall is changed (Handler et al, 2019; Aso and Rubin, 2016; Cohn et al, 2015; Burke et al, 2012). Evidence for this MBON balance model (Cognigni et al, 2018) has been provided for short-term memory (Aso et al, 2014b; Hige et al, 2015a; Perisse et al, 2016; Cohn et al, 2015) but to what extent, if any, such a model applies to long term memory remains unclear. Recently, genetic tools have been created that enable genetic tagging of long-term memory engram cells in the MB (Miyashita et al, 2018; Siegenthaler et al, 2019). These studies indicate that modulation of a relatively small number of KCs is sufficient to significantly alter memory recall.

Here, we generate a realistic computational model of the core KC>MBON module that is essential for the incorporation of memories to determine the cellular basis for decision making. Prior studies (Gouwens and Wilson, 2009) used “synthetic” (randomly generated) models of dendritic trees of *Drosophila* projection neurons to gain first insights into the computational processes of central neurons. Using linear cable theory, they demonstrated that voltage differences are much smaller within the dendritic trees than between the dendrites and the soma. Using a similar approach, Scheffer and coworkers simulated voltage distributions in the dendritic tuft of another *Drosophila* neuron (EPG) but using its measured morphology, rather than random geometries (Scheffer et al, 2020). Here, we go beyond these studies by combining precise structural data including the full synaptic connectivity from the electron microscopy-based synaptic connectome (Takemura et al, 2017) of the *Drosophila* MB with functional (electrophysiological) data of an individual MBON. This approach allows us to determine the impact of populations of KCs on MBON activity and to resolve the potential computational mechanisms underlying memory encoding. We focus on MBON-α3 (MBON-14 in the terminology of (Aso et al, 2014a)) an MBON at the tip of the α-lobe that is relevant for long term memory (Aso et al, 2014b; Plaçais et al, 2013). We first determine the physiological parameters of MBON-α3 by patch clamp electrophysiology. We then use these parameters to generate an anatomically and physiologically realistic *in silico* model of MBON-α3 and its KC inputs. Using a variety of computational simulations, we provide evidence that MBON-α3 is electrotonically compact and perfectly suited to incorporate a substantial number of memories based on largely stochastic KC inputs.

## 2 Results

### 2.1 Electrophysiological characterization of MBON-α3

MBON-α3 is particularly suitable for computational modeling due to the availability of its complete dendritic reconstruction through electron microscopy (Takemura et al, 2017). In addition, we can access MBON-α3 genetically by using a specific Gal4 line (MB082C-Gal4) to label MBON-α3 by the expression of membrane-bound GFP (10xUAS-IVS-mCD8::GFP) (Figure 1A). In immunohistochemical co-labelings with the active zone marker Bruchpilot (Brp), we were able to visualize the dendritic input sites, the axonal projections and the cell soma of MBON-α3 (Figure 1A, arrow). We used this labeling to perform fluorescence-guided electrophysiological whole-cell patch-clamp recordings of this neuron *ex vivo* (Figure 1C,D). We carried out standard patch-clamp protocols in five cells of independent preparations and recorded all essential physiological parameters (Table 1). It is of note that we cannot discriminate between the two different MBON-α3 cells that are marked by the split Gal4 line and that are almost identical with regard to their dendritic tree position and synaptic input number (Takemura et al, 2017). First, we performed a continuous current-clamp protocol over 60s at a sampling rate of 50kHz which allowed us to determine the average resting membrane potential as −56.7 ± 2.0mV. This value for the resting membrane potential is in line with prior measurements of MBONs *in vivo* and of other central neurons in *Drosophila* (Hige et al, 2015a; Wilson, 2004; Gu and O’Dowd, 2006; Groschner et al, 2022). In these recordings, we observed spontaneous firing activity of MBON-α3 with an average frequency of 12.1Hz. Next, we performed a current-clamp de- and hyper-polarization protocol where we recorded 400ms trials at a sampling rate of 50kHz. Each trial started with 0pA for 10ms, followed by a current injection of −26pA to 32pA, with 2pA increments for 400ms. These protocols allowed us to evoke action potentials and to determine the resulting changes in membrane potential. An example displaying selected traces of a representative recording is shown in Figure 1F. Two additional examples are illustrated in Figure 1 Supplement 1. From this data we determined the average membrane time constant as *τ*_m_ of 16.06 ± 2.3ms. We then performed a voltage-clamp step protocol without compensating for the series resistance. In this experiment, we recorded at 20kHz while applying a 5mV voltage step for 100ms. This allowed us to calculate the average membrane resistance as R_m_ = 926 55M*Ω*, a rather high membrane resistance when compared to similarly sized neurons (Gouwens and Wilson, 2009). MBON-α3 is thus highly excitable, with small input currents being sufficient to significantly alter its membrane potential (Gouwens and Wilson, 2009; Sakmann and Neher, 1995). The amplitude of action potentials was rather small (4.3 ± 0.029ms) but similar to previously reported values for different MBONs (Hige et al, 2015a). The small amplitudes are likely a result of the unipolar morphology of the neuron with a long neurite connecting the dendritic input region to the soma (see below). As a result, signal propagation may partially bypass the soma (Figure 1F) (Gouwens and Wilson, 2009). We calculated the neuron’s membrane capacitance (C_m_ = tau°Rm) as C_m_ = 16.76 ± 1.90pF, which classifies MBON-α3 as a mid-sized neuron (Gouwens and Wilson, 2009).

**Table 1.**
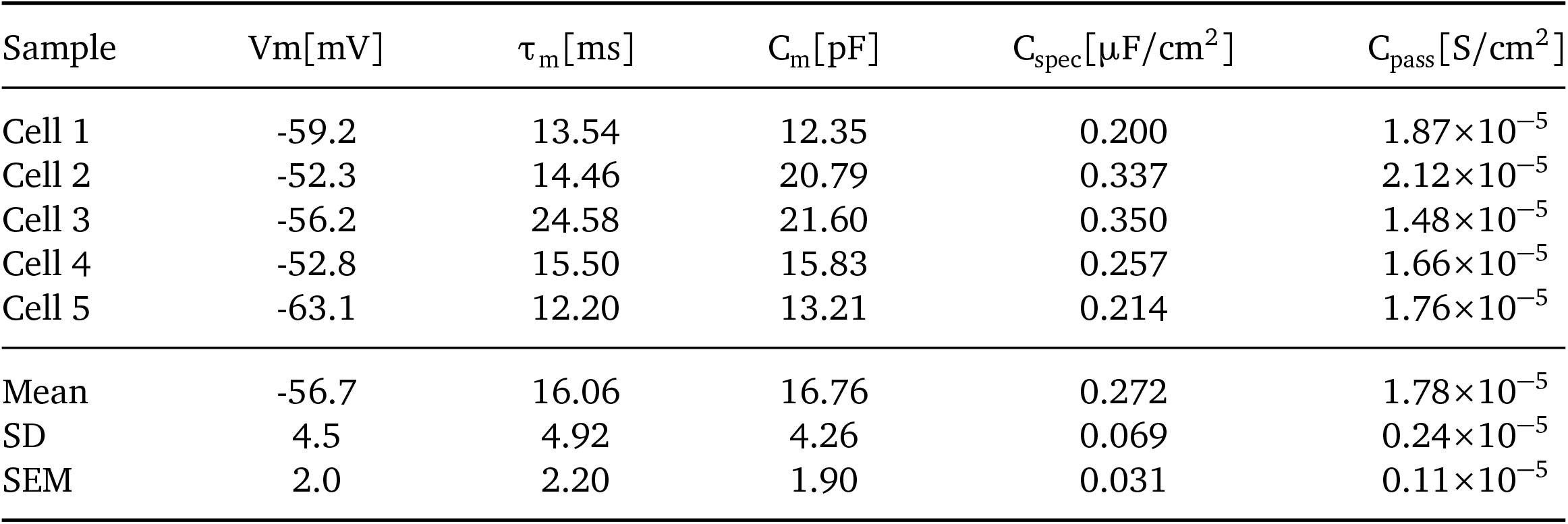
Summary of passive membrane properties of MBON-α3, calculated from electrophysiological measurements from five neurons. From left to right: resting membrane potential (V_m_) in mV, membrane time constant (*τ*_m_) in ms, capacitance of the membrane (C_m_) in pF, specific capacitance (C_spec_) in *μ*F°cm^2^ and passive membrane conductance (C_pass_) in S°cm^2^. **Table 1 Source Data 1** Summary of passive membrane properties of MBON-α3, calculated from electrophysiological measurements from four neurons marked by cytoplasmic EGFP.

| Sample | $V_m[\text{mV}]$ | $\tau_m[\text{ms}]$ | $C_m[\text{pF}]$ | $C_{\text{spec}}[\mu\text{F}/\text{cm}^2]$ | $C_{\text{pass}}[\text{S}/\text{cm}^2]$ |
| --- | --- | --- | --- | --- | --- |
| Cell 1 | -59.2 | 13.54 | 12.35 | 0.200 | $1.87 \times 10^{-5}$ |
| Cell 2 | -52.3 | 14.46 | 20.79 | 0.337 | $2.12 \times 10^{-5}$ |
| Cell 3 | -56.2 | 24.58 | 21.60 | 0.350 | $1.48 \times 10^{-5}$ |
| Cell 4 | -52.8 | 15.50 | 15.83 | 0.257 | $1.66 \times 10^{-5}$ |
| Cell 5 | -63.1 | 12.20 | 13.21 | 0.214 | $1.76 \times 10^{-5}$ |
| Mean | -56.7 | 16.06 | 16.76 | 0.272 | $1.78 \times 10^{-5}$ |
| SD | 4.5 | 4.92 | 4.26 | 0.069 | $0.24 \times 10^{-5}$ |
| SEM | 2.0 | 2.20 | 1.90 | 0.031 | $0.11 \times 10^{-5}$ |

**Figure 1.**
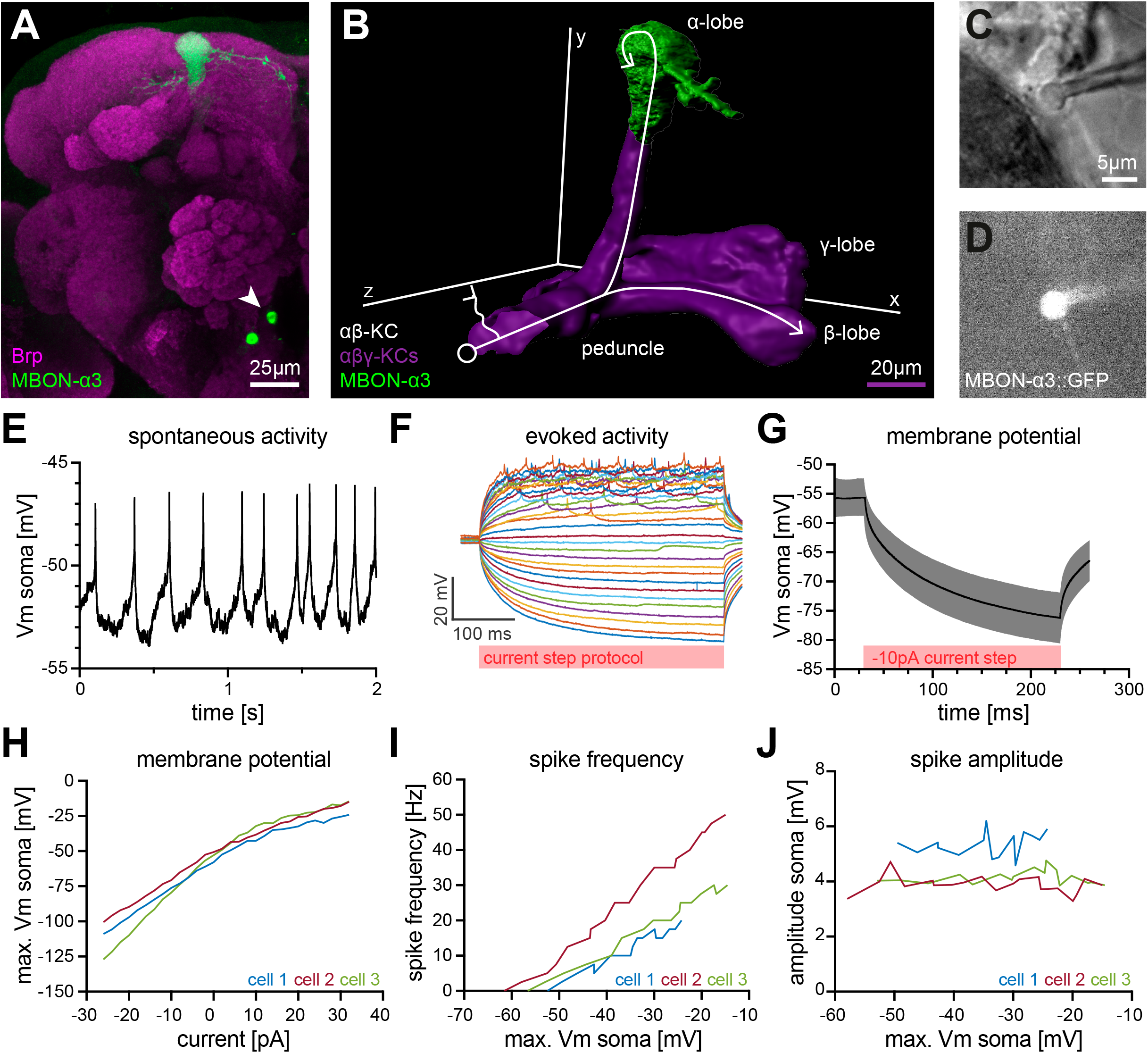
Electrophysiological characterization of the MBON-α3 neuron. (A) Immunohistochemical analysis of the MBON-α3 (green, marked by GFP expression) reveals the unique morphology of the MBON with its dendritic tree at the tip of the α-lobe and the cell soma (arrowhead) at a ventromedial position below the antennal lobes. The brain architecture is revealed by a co-staining with the synaptic active zone marker Bruchpilot (Brp, magenta). (B) Artificial superimposition of a partial rendering of MBON-α3 (green) and of *αβγ*-KCs (magenta) to illustrate KC>MBON connectivity. A simplified schematic *αβ*-KC projection is illustrated in white. (C) Transmitted light image of an MBON-α3 cell body attached to a patch pipette. (D) Visualisation of the GFP-expression of MBON-α3 that was used to identify the cell via fluorescence microscopy. The patch pipette tip is attached to the soma (right side). The scale bar in C applies to C and D. (E) Recording of spontaneous activity of an MBON-α3 neuron in current clamp without current injection. (F) Recording of evoked neuronal activity of an MBON-α3 neuron with step-wise increasing injection of 400ms current pulses. Pulses start at −26pA (bottom) with increasing 2pA steps and end at +32pA (top). (G) Mean trace of the induced membrane polarization resulting from a 200ms long current injection of −10pA. 105 trials from three different cells (35 each) were averaged. Gray shading indicates the SD. (H) Relative change of membrane potential after current injection. Vm is equal to the maximum depolarization of the membrane. (I) Correlation between the frequency of action potential firing and changes in membrane potential. (J) Relative change of spike amplitude after current injections. Spike amplitude represents the peak of the largest action potential minus baseline. Also see **Figure 1 Supplement 1** and **Figure 1 Supplement 2**.

To enable the calibration of the *in silico* model, we then recorded 35 trials per cell in a current-clamp injection protocol with injections of 10pA current pulses for 200ms at a sampling rate of 20kHz. We averaged 105 traces from 3 selected example cells to a standard curve, describing the membrane kinetics of MBON-α3 in response to current injections (Figure 1G). To further characterize MBON-α3, we analyzed the resulting data from the current-clamp de- and hyper-polarization protocols in the three example cells (Figure 1F and Figure 1 Supplement 1). We recorded the absolute deflection, the action potential firing frequency, and the amplitude of action potentials for all individual cells. In all three cells, we observed that the absolute deflection of the membrane potential was proportional to the increasing current injections (Figure 1H). The action potential firing frequency increased gradually with increasing membrane potential deflections (Figure 1I), with no significant change in action potential amplitudes (Figure 1J). The same effect was observed with an increasing current injection (Figure 1 Supplement 1). This is consistent with the general idea that action potentials are “all or nothing” events, and that MBON-α3 is a spike-frequency adapting neuron constistent with prior observations of other MBONs (Hige et al, 2015a). We repeated these recordings with a cytosolic EGFP as an independent and non-membrane associated marker and observed consistent results (Figure 1 Supplement 2 and Table 1 Source Data 1). We next used these data to generate an *in silico* model based on realistic passive membrane properties.

### 2.2 Constructing an *in silico* model of MBON-α3

For our *in silico* model of MBON-α3, we used the recently published electron microscopy-based connectome of the *Drosophila* mushroom body (Takemura et al, 2017). This dataset includes the precise morphological parameters of the entire dendritic tree including the precise location data for all 12,770 synaptic connections from the 948 innervating KCs. It is important to note that this dataset represents a single fly and the EM-based reconstruction was not able to faithfully assign all synaptic connections (potentially missing 10% of synapses) (Takemura et al, 2017). The comparison with the hemibrain data set (Scheffer et al, 2020) that provides a second independent reconstruction of the MBON from a different fly showed very few differences in KC to MBON-α3 connectivity. Thus, despite potential shortcomings, this dataset represents the most accurate template to reconstruct the dendritic tree and connectivity of an MBON. To build a complete morphological reconstruction of MBON-α3, we included the axonal reconstructions from the hemibrain dataset (Scheffer et al, 2020) and determined the length of the soma to the dendrite via confocal light microscopy. We incorporated these data together with all other values into the NEURON simulation environment (Hines and Carnevale, 1997) (Figure 2 A & C and Figure 2 Supplement 1).

**Figure 2.**
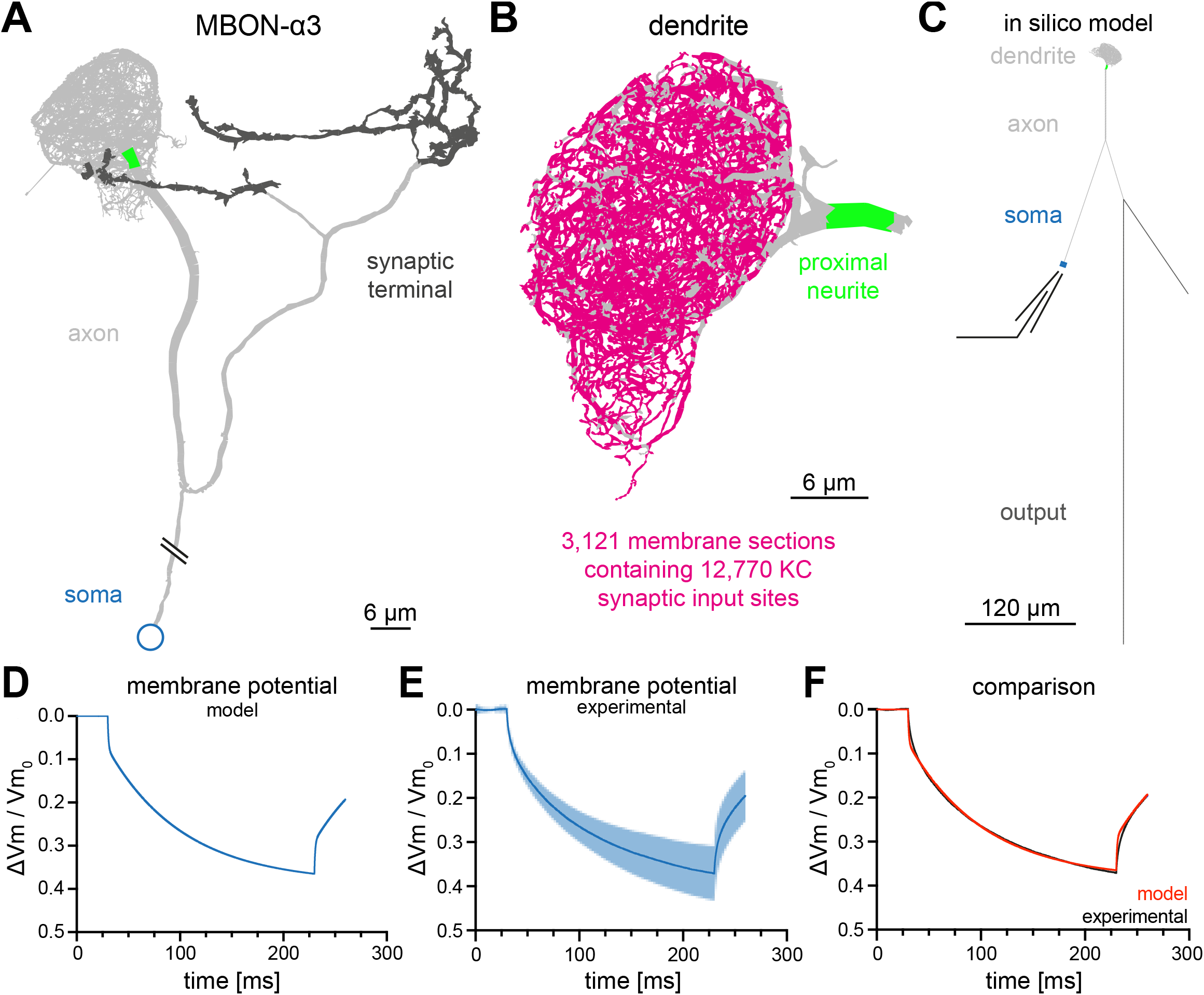
Construction of a computational model of MBON-α3. (A) Electron microscopy based reconstruction of MBON-α3. Data was obtained from NeuPrint (Takemura et al, 2017) and visualized with NEURON (Hines and Carnevale, 1997). Data set MBON14 (ID 54977) was used for the dendritic architecture (A-C) and MBON14(a3)R (ID 300972942) was used to model the axon (light gray) and synaptic terminal (dark gray, A, C). The proximal neurite (bright green) was defined as the proximal axonal region next to the dendritic tree. This is the presumed site of action potential generation. Connectivity to the soma (blue) is included for illustration of the overall morphology and is not drawn to scale. (B) A magnified side view of the dendritic tree of MBON-α3. Within the dendritic tree a total of 4,336 individual membrane sections were defined (light gray area). The 3,121 membrane sections that contain one or more of the 12,770 synaptic input sites from the 948 innervating KCs are highlighted in magenta. (C) Simplified *in silico* model of MBON-α3 highlighting the size of the individual neuronal segments. We simulated recordings at the proximal neurite (specified as the potential site of action potential generation based on morphological parameters; this area is still included in the dendritic tree reconstruction of Takemura et al. 2017) or at the soma. Values for the different sections are provided in **Figure 2 Supplement 1, Figure 2 Source Data 1**. (D) Normalized trace of a simulated membrane polarization after injection of a 200ms long square-pulse current of − 10pA at the soma. (E) Normalized and averaged experimental traces (blue) with standard deviation (light blue) of our measured depolarization. (F) Comparison of a normalized induced depolarization from the model (red) and from the experimental approach (blue). The model was fitted within the NEURON environment to the measured normalized mean traces with a mean squared error between model and measured data of 0.048mV^2^. Also see **Figure 2 Supplement 1, Figure 2 Supplement 2** and **Figure 2 Source Data 1**.

Based on the published vector data (Takemura et al, 2017) we subdivided the dendritic tree into 4,336 linear sections, of which 3,121 are postsynaptic to one or more of the 12,770 synaptic contacts from 948 KCs (Figure 2B). We used linear cable theory (Niebur, 2008) to generate the *in silico* model of MBON-α3. Linear cable theory is the baseline model for neurites and the most rational approach in the absence of detailed information about potential nonlinear currents.

In previous simulations of *Drosophila* neurons, the kinetics of membrane polarisation as a response to current injections were adjusted to *in vivo* measurements through a fitting procedure (Gouwens and Wilson, 2009). We first defined the boundaries of the passive membrane parameters within NEURON’s PRAXIS (principle axis) optimization algorithm and incorporated the experimentally defined passive membrane properties (Table 1).

We then determined the biophysical parameters of the *in silico* neuron by current step injections of −10pA for 200ms at the soma and by recording the neuronal voltage traces. This enabled us to fit the membrane parameters of our model neuron to the experimental data of the three example cells (Figure 2E). The biophysical parameters that resulted from the fitting procedure are provided in Table 2.

**Table 2.**
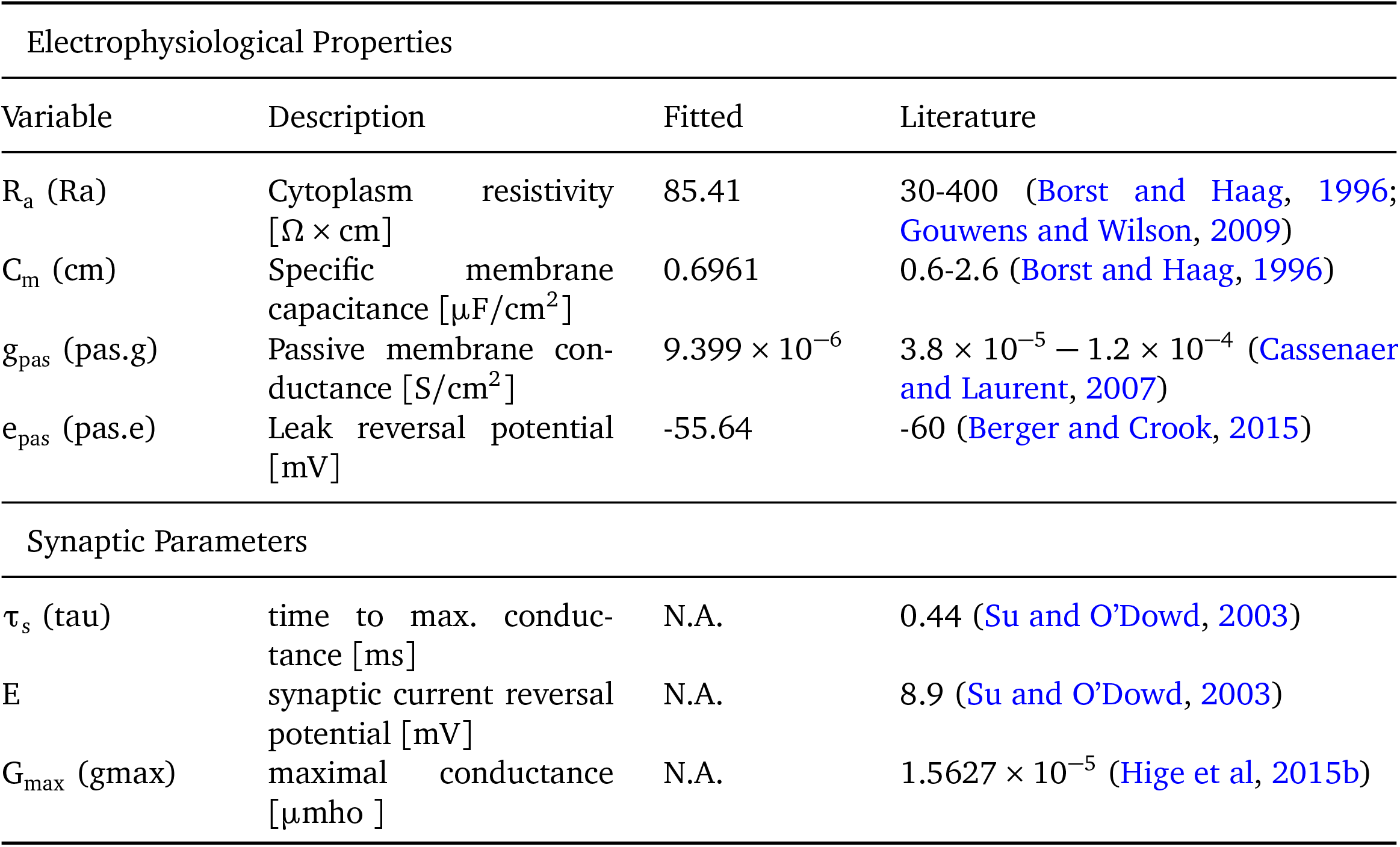
Electrophysiological values applied for the *in silico* model of MBON-α3-A. Parameter values were obtained from literature or were based on the fitting to our electrophysiological data. The first column shows the parameters, with the names used in the NEURON environment in parentheses. The maximal conductance was determined to achieve the target MBON depolarization from monosynaptic KC innervation.

We obtained membrane kinetics that closely resemble *ex vivo* current injections of −10pA, with a mean squared error between model and measured data of 0.0481mV^2^. For comparison, we normalized both the modelled current injection (Figure 2D) and the average of the traces from the *ex vivo* recordings (Figure 2E) and overlayed the resulting traces in Figure 2F.

Note that the faster voltage change at the onset and offset of the current injection seen in the simulation is caused by the small size of the soma which results in a much faster “local” somatic time constant than that of the whole cell (Figure 2 Supplement 2). This is also visible in the average of the recorded traces (Fig. 2F), but more obvious in individual (not trial-averaged) recorded traces (Figure 2 Supplement 2C). Overall, these results indicate that we were able to generate a realistic *in silico* model of MBON-α3.

### 2.3 Effect of KC>MBON synaptic inputs

As KCs are cholinergic (Barnstedt et al, 2016) we utilized previously determined parameters of cholinergic synapses to simulate synaptic input to MBON-α3. We used a reversal potential of 8.9mV and a synaptic time constant *τ*_s_ = 0.44ms as target values (Su and O’Dowd, 2003). Individual synaptic contacts were simulated as alpha functions (g_max_(t − t_i_) × exp(− (t −t_i_ *−τ*_s_)°*τ*_s_)) where t_i_ is the time of the incoming spike and g_max_ = 1.56 × 10^−11^S, chosen to obtain response levels in agreement with the target values. The average number of synaptic contacts from a KC to this MBON is 13.47 (Takemura et al, 2017), Fig. 4B. To determine the computational constraints of MBON-α3, we first stimulated each of the 3,121 dendritic sections that receive KC synaptic innervation using the described alpha-function conductance change. We then “recorded” (from our simulation) the resulting voltage excursions at the dendritic input site (Fig 3A, red), at the proximal neurite (putative action potential initiation site, green) and at the cell soma (blue). The comparative analysis of the responses at these three locations revealed two notable features: (1) voltage excursions directly within the dendrite are faster and substantially larger than in the proximal neurite and soma, and (2), the resulting voltages in the latter two compartments are much less variable than in the dendrite and distributed within a very small voltage range (Figure 3D,E).

**Figure 3.**
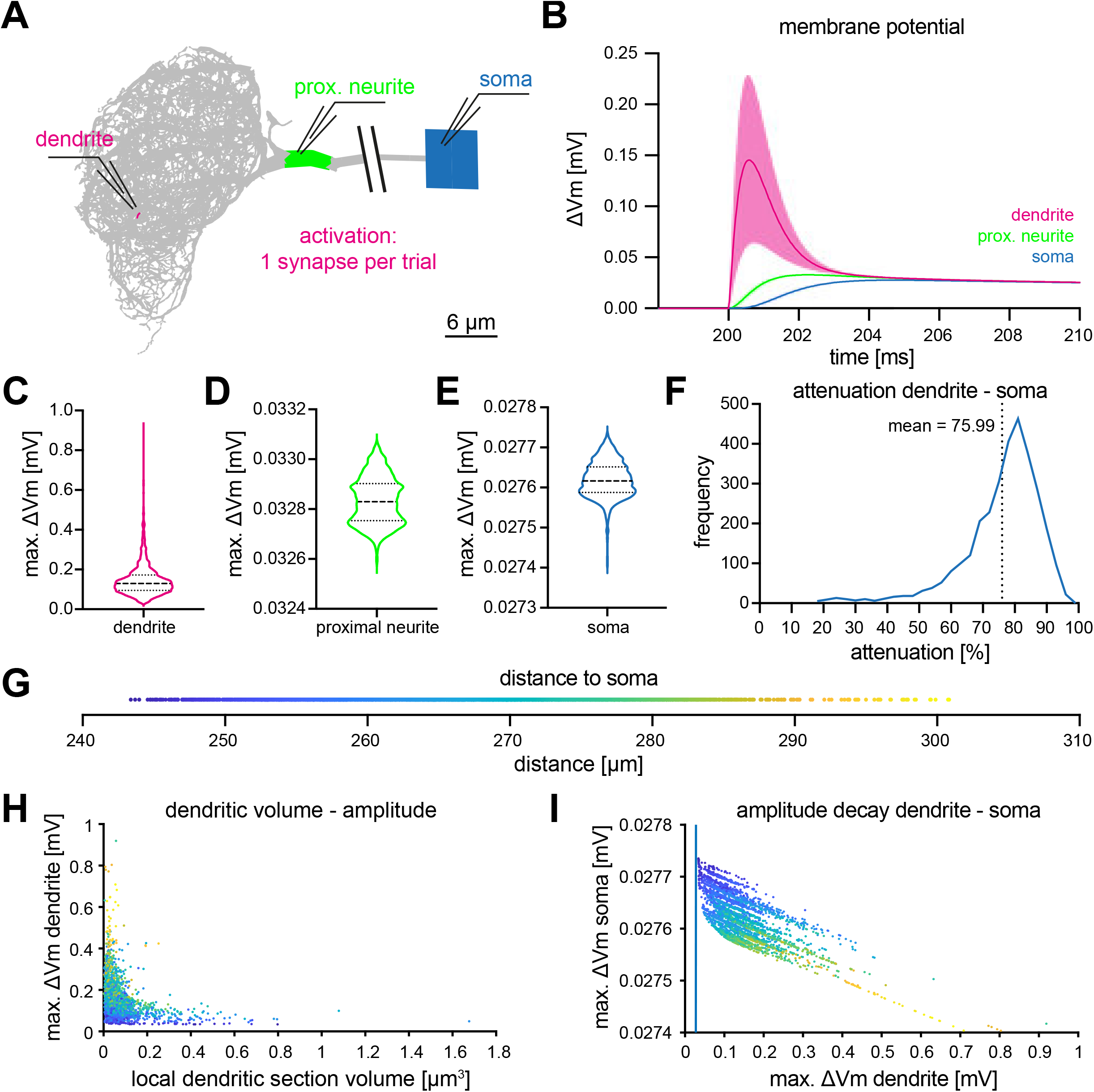
Computational characterization of MBON-α3. (A) Simulated stimulation of individual synaptic input sites. Voltage deflections of synaptic currents were analyzed locally at the dendritic segment (red, next to electrode tip), the proximal neurite (green) and the soma (blue). The soma is depicted as a square that reflects its size in the NEURON environment. Color code also applies to B-F. (B) Resulting mean voltages (lines) and standard deviations (shading) from synaptic activations in 3,121 individual stimulus locations (dendrite), and at the proximal neurite and soma. (C-E) Violin plots of the simulated maximal amplitudes at the dendritic section (C), the proximal neurite (D) and the soma (E) in response to synaptic activations. Dotted lines represents the quartiles and dashed lines the mean. Proportions of the violin body represent the distribution of individual data points. Note the difference in scale and compactification of the amplitudes in the proximal neurite and soma. (F) Distribution of the percentage of voltage attenuation recorded at the soma. (G) Distribution of distance between all individual dendritic sections and the soma. This color code is applied in H and I. (H) The elicited local dendritic depolarization is plotted as a function of the local dendritic section volume. (I) The soma amplitude is plotted as a function of the amplitude at individual dendritic segments. Note different scales between abscissa and ordinate; the blue line represents identity. Also see **Figure 3 Supplement 1**.

**Figure 4.**
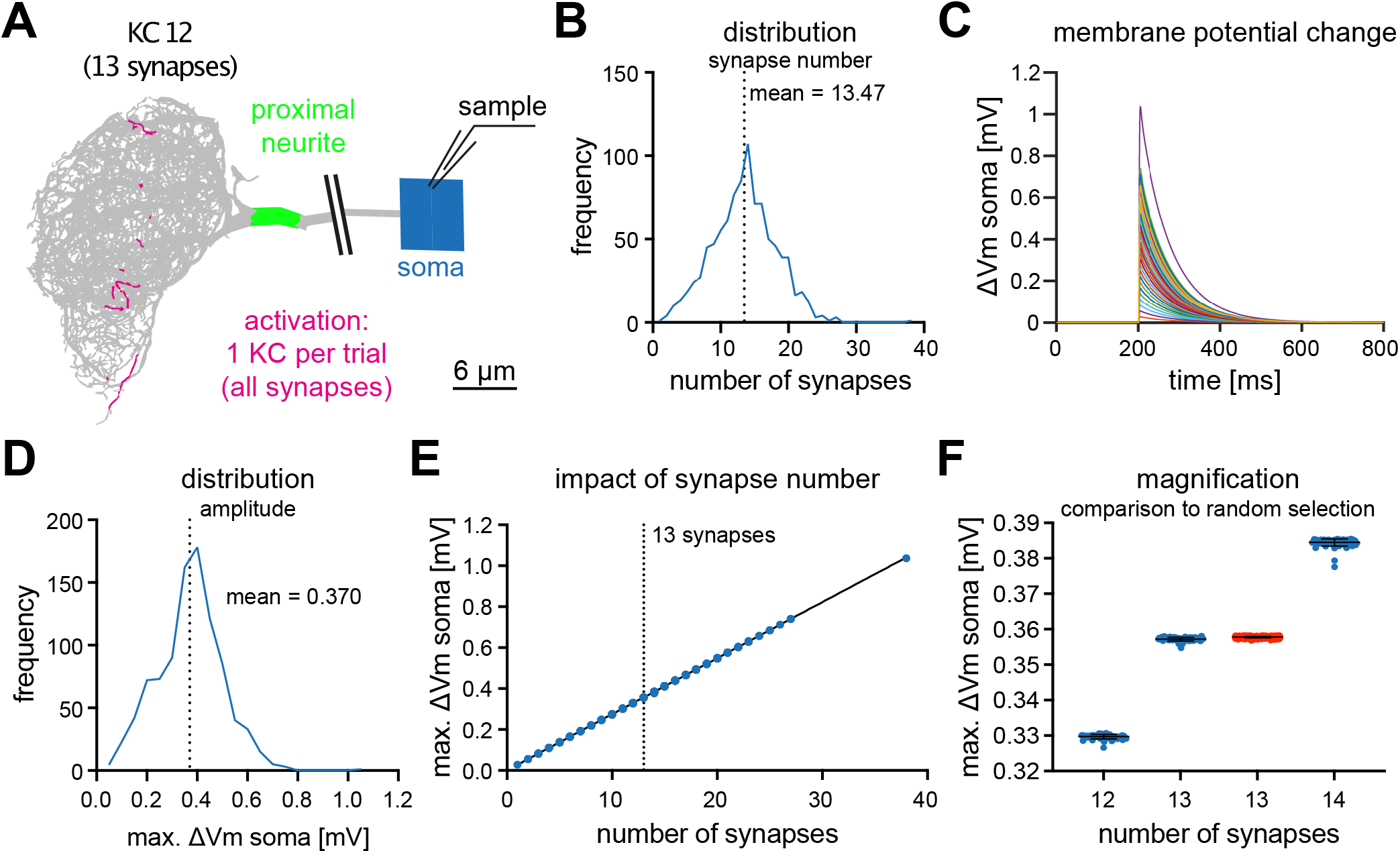
MBON-α3 is electrotonically compact. (A) Simulated activation of all synapses from all 948 individual KC innervating MBON-α3. A representative example of a KC (KC12) with 13 input synapses to MBON-α3 is shown. (B) Histogram of the distribution of the number of synapses per KC. (C) Membrane potential traces from simulated activation of all 948 KCs. (D) Distribution of the elicited amplitudes evoked by the individual activations of all KCs. (E) Correlation between the somatic amplitude and the number of activated synapses in the trials. (F) Blue: Distribution of the somatic voltage change evoked by the activation of all KCs with exactly 12, 13, or 14 input synapses. Each dot represents one simulated KC. Red: Same, but for activation of 13 random synapses per simulation in 1000 independent trials.

These results indicate that the morphological architecture and biophysical parameters of MBON-α3 promote a “compactification,” or “democratization,” of synaptic inputs, with all inputs resulting in similar voltage excursions at the proximal neurite and soma, regardless of the strength of the initial dendritic voltage or the position on the dendritic tree.

To a first approximation, compactification can be understood from a simple analytical estimate of the space constant of this neuron. It is obtained by assuming a non-branching cylinder with the mean diameter of the dendritic segments, 0.29*μ*m, as obtained from the morphological data from Takemura et al (2017). Taking the values for the conductance of the cytoplasm of (r_a_ = 85*Ω*cm) and the transmembrane conductance (g_pas_ = 9.4 × 10^−6^S°cm^2^) from Table 2, the characteristic length over which excitations decay can then be computed by linear cable theory (Niebur, 2008) as *λ* ≈ 1, 300*μ*m. This is about twice the maximal path length between any two segments of the cell, see Fig. 2C. Using the average diameter of all neuronal compartments, both in the dendritic tree and in the other components of the neuron as listed in Fig 1 Supplement 1, increases *λ* by about 10% because the non-dendritic segments have larger diameters. In either case, synaptic input even at distal locations on the dendritic tree can influence somatic voltage deflections nearly as strongly as more proximal synapses, see Figure 3E.

To understand additional influences of cell morphology for the input computation, we analyzed the relation between synaptic position and local dendritic volume and the resulting voltages in the dendrite, proximal neurite and soma. Representing the distance between dendritic inputs and soma (Figure 3G) by a color code, we find that as in many tree-like structures, there is a systematic tendency that more distal segments (more distant from the soma) have smaller volume and diameter, Figure 3H. The same figure also shows that the simulated voltage excursions are many times larger in the distal, small segments than in proximal, large segments, over a range from ≈0.03mV to ≈ 1mV. This is expected: placing an electrical charge (integrated synaptic current) on a small capacitor (low-volume segment with small surface area) results in a larger voltage excursion than placing the same charge on a large capacitor (large segment). It is highly remarkable, however, that the morphology of the MBON-α-3 dendritic tree seems finely tuned, with the goal of maximizing the equality of the functional weight of all synapses. Specifically, the decrease of the voltage excursion in the soma caused by the injection of a current is nearly perfectly (within less than 0.5*μ*V) compensated by the increase of the local dendritic voltage due to the smaller volume of distal segments, see Figure 3I. The approximately 30-fold difference of local dendritic voltages (≈0.03mV to ≈ 1mV) is nearly exactly compensated, such that activation of all synapses having exactly the same effect at the soma, within less than one *μ*V.

We next analyzed the impact of individual KCs on MBON-α3. We simulated the activation of all synapses of each individual KC that innervates MBON-α3, using the synaptic parameters described above, and we recorded the simulated voltage excursions in the soma, Figure 4A. The number of synapses varied between KCs, ranging from 1 to 27 (plus one outlier with 38 synaptic contacts), with a mean of 13.47 synapses per KC (Figure 4B).

In these simulations, we observed a wide range of voltage excursions (note the outlier in Figure 4B,C,D,E). The neuron is firmly in a small-signal operation mode, as we observed a highly linear relation between the voltage excursion and the number of synapses per KC, Fig 4E. Activating a single KC leads to a voltage excursion at the soma with a mean of 0.37mV, Fig 4D. To further analyze this linear relationship between synapse number and somatic voltage excursion, we analyzed all KCs with exactly 12, 13 or 14 synaptic inputs to MBON-α3. This analysis revealed a highly stereotypical depolarization of the soma for all these KCs, despite wide variations in the position of the individual synaptic contacts (Figure 4E,F). Activation of 13 random synapses in 1000 independent trials resulted in similar voltage excursions. Interestingly, in these simulations the variance of the responses was smaller compared to the activation of the individual KCs (Figure 4F). Together, these results provide further support for the electrotonical compactness of MBON-α3.

### 2.4 Physiological and tuned activation of MBON-α3

A given odor activates only a small fraction of KCs reliably, reported as about 5% in some studies (Honegger et al, 2011; Siegenthaler et al, 2019) and 6% in others (Turner et al, 2008; Campbell et al, 2013). This one-percentage point difference may well be caused by differences in the experimental preparations. However, in absolute terms a change from 5% to 6% in the *same* system represents a 20% increase. To understand the physiological effect of differences in the number of activated KCs of this size, we first established a baseline condition by simulating voltage excursions in the soma resulting from the simultaneous activation of sets of 50 KCs (corresponding to 5%), randomly selected from the 984 KCs innervating MBON-α3.

In 1,000 independent activation trials (Figure 5A), we observed highly stereotypical activation patterns, with a mean somatic depolarization of 15.24mV and only small variations in the time course (Figure 5B,C (blue)). These results are consistent with an activation of 50 KCs being sufficient to induce changes in the firing frequency of MBON-α3 (Figure 1F). The fact that all simulations resulted in a significant depolarization consistent with action potential generation or alterations of the firing frequency together with the small range of variation within this dataset, provides support for the current model that odor encoding in KCs is, at least to a large extent, random (stochastic) (Caron et al, 2013; Zheng et al, 2022; Li et al, 2020; Eichler et al, 2017).

**Figure 5.**
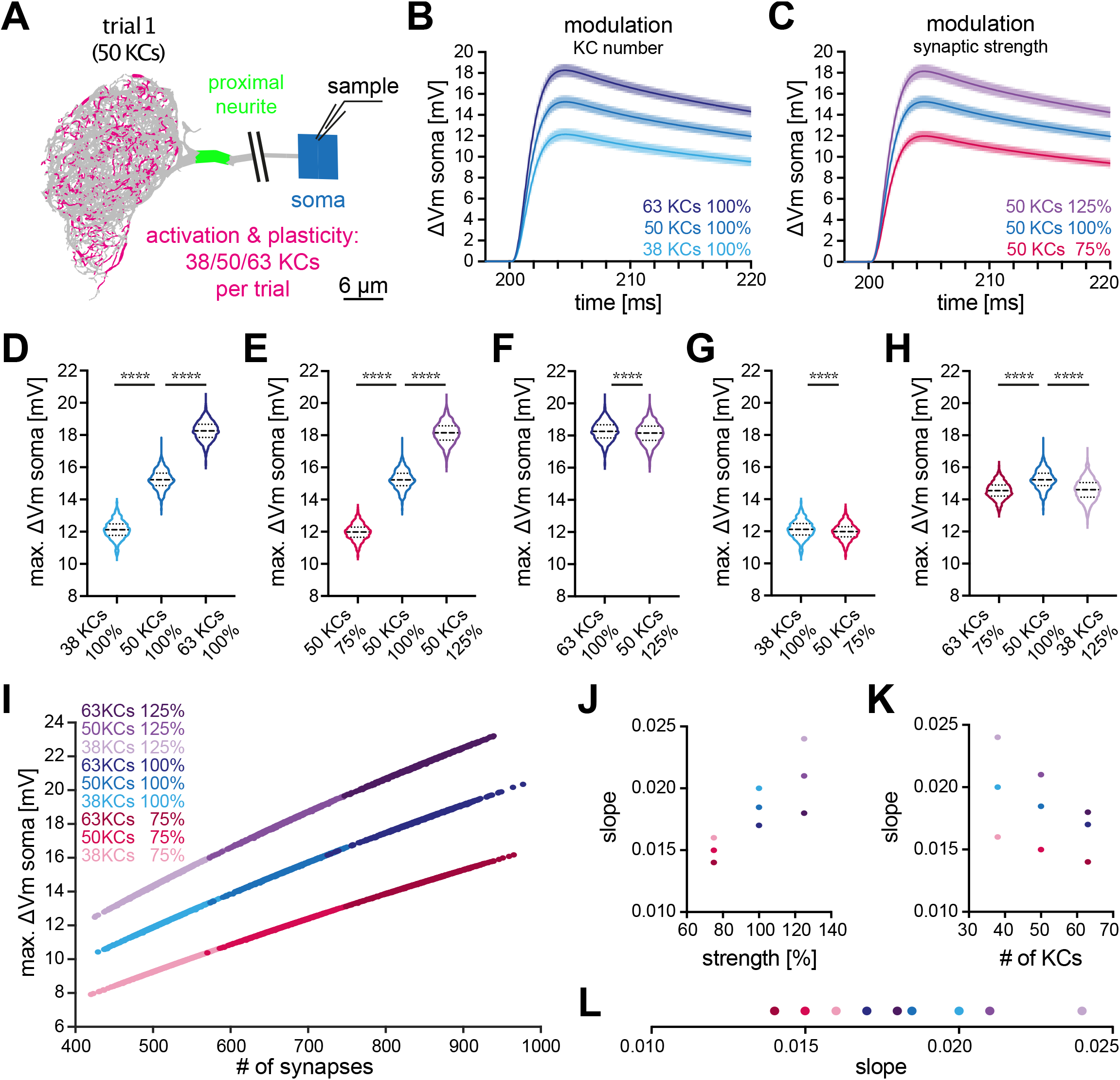
Effect of KC recruitment and synaptic plasticity on MBON-α3 responses. (A) Schematic of the simulated recording paradigm. We modeled the activation of ≈5% of KCs of random combinations of identified KCs, *i*.*e*. we activated the anatomically correct synapse locations of 50 randomly selected KCs to mimic activation by an odor. The panel shows the distribution of activated synapses for one of these trials. We then varied the number of activated KCs by ±25% around its mean of 50, *i*.*e*. 38, 50 and 63 KCs, and we varied synaptic strength ±25% around its original value, *i*.*e*. 75%, 100% and 125%. (B) Mean somatic depolarizations after activating 1000 different sets of either 38 (light blue), 50 (blue), or 63 (dark blue) KCs, all at standard synaptic strength (100%). Shades represent the standard deviations. (C) Mean somatic depolarizations after activating 1000 different sets of 50 KCs at either 75% (red), 100% (blue) or 125% (purple) synaptic strength. Shades represent the standard deviations. (D, E) Violin plots of the relative amplitudes evoked by the different KC sets from (B, C). (F-G) Violin plots for the comparison of the amplitudes between the different plasticity paradigms. (F) 63 KCs at 100% *vs*. 50 KCs at 125% synaptic strength. (G) 38 KCs at 100% *vs*. 50 KCs at 75% synaptic strength. Plots in D-G use the color code from (B) and (C). (H) 63 KCs at 75% (dark red) *vs*. 50 KCs at 100% (blue) and 38 KCs at 125% (light purple) synaptic strength. See Fig. 3 for symbols in violin plots. (I) Relation between somatic depolarization and the number and strength of activated synapses. Illustrated are all tested conditions for synaptic plasticity and KC recruitment. Stimulation paradigms are color coded as shown. (J) Relation between the slope of the condition-specific linear regressions from (I) and the synaptic strength of the activated KC sets. (K) Relation between the slope of the condition-specific linear regressions from (I) and the number of recruited KCs. (L) Slopes of the condition-specific linear regressions from (I). Color codes of (J-L) as in (I). Statistical significance was tested in multiple comparisons with a parametric one-way ANOVA test (D, E and H) resulting in p=<0.0001 for all conditions, or in an unpaired parametric t-test (F and G) resulting in p=<0.0001 for all conditions. Numerical results are summarized in Table 3.

Based on *in vivo* data, two different modes of plasticity may act at the level of the KC-MBON module to alter MBON output to conditioned odors. *In vivo* imaging of short-term memory formation demonstrated alterations of KC input strength (Owald and Waddell, 2015; Hige et al, 2015a; Perisse et al, 2016; Plaçais et al, 2013; Pai et al, 2013; Séjourné et al, 2011; Jacob and Waddell, 2020) while recent observations of long-term memory demonstrated a specific change (increase) in the number of KCs representing the conditioned odor (Delestro et al, 2020; Shyu et al, 2019; Baltruschat et al, 2021). To directly address the relative impact of these two modulations on MBON depolarization, we repeated the same simulation but now modified input strength by either altering activated KC numbers or by modulating the synaptic gain. To enable a direct comparison, we modulated each factor by 25%. First, we increased the number of activated KCs per set by 25%, from 50 to 63. This lead to a mean somatic depolarization of 18mV, a significant increase by 19% (Figure 5B,D; dark blue). Decreasing the number of activated KCs by 25%, from 50 to 38, lead to a mean somatic depolarization of 12mV, a significant decrease by 20% (Figure 5B,D; light blue). Next, we either increased or decreased synaptic strength within sets by 25%. Increase lead to a mean somatic depolarization of 18mV, a significant increase by 19% (Figure 5C,E; purple) while decrease lead to a mean somatic depolarization of 12mV, a significant decrease by 21% (Figure 5C,E; red). Averaged over all conditions, a change of input of 25% lead to a change of 20±1% in membrane depolarization.

**Table 3.**
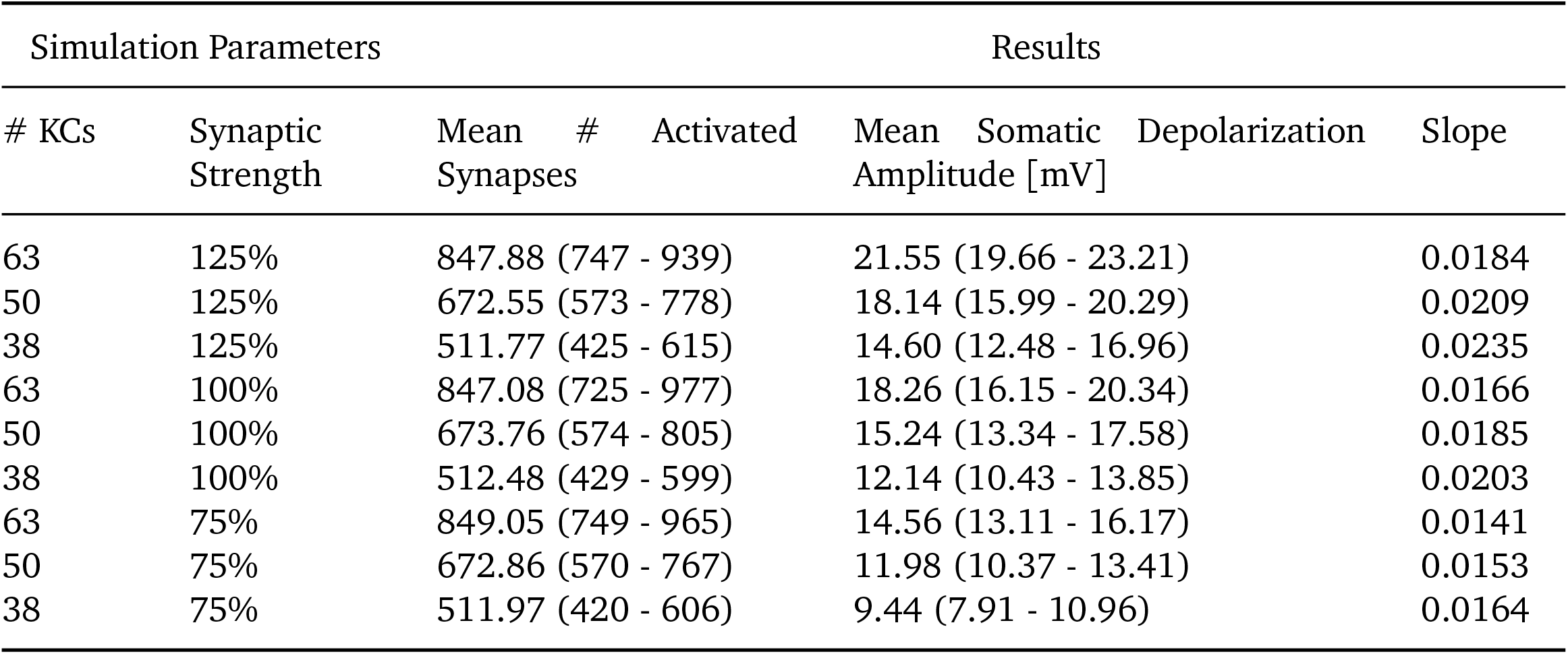
Summary of data from the plasticity simulations of Figure 5. Simulations vary the number of active KCs and/or the strength of the KC-MBON synapses by scaling the maximal conductance parameter of the synaptic alpha-conductance function. Results include the mean number of activated synaptic sites as well as the mean amplitude of the somatic voltage depolarization across the 1000 trial repetitions of each simulation. Ranges are reported as parentheticals. The slope of the linear regression describing the relation between the number of active synapses and somatic voltage amplitude is reported (Fig. 5I-L).

Even though the integrated synaptic conductance was changed by the same amount by our manipulation of either synaptic strength or number of activated KCs, we observed small but significant differences in the resultant mean somatic amplitudes (Figure 5F,G). Finally, we combined both manipulations by compensating an increase in KC number with a corresponding decrease in synaptic strength and vice versa. An overview of the performed simulations and numerical results are provided in Table 3.

Both manipulations resulted, on average, in a significant decrease of MBON membrane depolarization (Figure 5H). Our simulations revealed a nearly linear relationship between the activated number of synapses and the measured amplitude in the soma regardless of synaptic strength (Figure 5I). Interestingly, we observed some differences in the slope of the responses between the different tuning modalities (Figure 5J,K,L), indicating that these modulations are not completely equivalent.

## 3 Discussion

For our understanding of the computational principles underlying learning and memory, it is essential to determine the intrinsic contributions of neuronal circuit architecture. Associative olfactory memory formation in *Drosophila* provides an excellent model system to investigate such circuit motifs, as a large number of different odors can be associated with either approach or rejection behavior through the formation of both short- and long-term memory within the MB circuitry (Aso and Rubin, 2016). In contrast to most axon guidance processes in *Drosophila* that are essentially identical in all wild-type individuals, processing of odor information in the MB largely, but not exclusively, depends on the stochastic connectivity of projection neurons to KCs that relay odor information from the olfactory glomeruli to the MBONs (Caron et al, 2013; Gruntman and Turner, 2013; Litwin-Kumar et al, 2017; Warth Pérez Arias et al, 2020; Zheng et al, 2022). However, the role of MBONs within the circuit is largely fixed between animals and many MBONs can be classified as either approach or avoidance neurons in specific behavioral paradigms (Aso et al, 2014b,a; Litwin-Kumar et al, 2017; Li et al, 2020; Takemura et al, 2017; Scheffer et al, 2020). Furthermore, individual KCs are not biased in their MBON connectivity but innervate both kinds of output modules, for both approach and avoidance. Learning and memory, the modulation of odor response behavior through pairing of an individual odor with either positive or negative valence, is incorporated *via* the local activity of valence-encoding DANs that can either depress or potentiate KC>MBON synaptic connections (Aso and Rubin, 2016; Berry et al, 2018; Handler et al, 2019; Yamagata et al, 2015; Cohn et al, 2015; Burke et al, 2012; Hige et al, 2015a; Owald et al, 2015a; Jacob and Waddell, 2020; Pai et al, 2013; Plaçais et al, 2013; Séjourné et al, 2011; Bouzaiane et al, 2015). As KC odor specificity and connectivity differ significantly between flies with respect to the number and position of KC>MBON synapses, this circuit module must be based on architectural features supporting robust formation of multiple memories regardless of specific individual KC connectivity.

Prior computational work addressing properties of central nervous system neurons in *Drosophila* relied on synthetic (randomly generated) data (Gouwens and Wilson, 2009) or partial neuronal reconstructions (Scheffer et al, 2020; Tobin et al, 2017). We go beyond these previous studies by combining precise structural data from the electron-microscopy based synaptic connectome (Takemura et al, 2017) with functional (electrophysiological) data. The structural data consists of the neuroanatomical structure of MBON-α3, including the 12,770 synaptic inputs of all its 948 innervating KCs. While this EM reconstruction may contain some mistakes in synaptic connectivity as e.g. up to 10% of synaptic sites remained unassigned within the dataset (Takemura et al, 2017), it currently represents the best possible template for an *in silico* reconstruction. The neuron’s functional properties were determined from *ex vivo* patch clamp recordings. Near-perfect agreement between experimentally observed and simulated voltage traces recorded in the soma shows that linear cable theory is an excellent model for information integration in this system.

Together, we obtain a realistic *in silico* model of a central computational module of memory-modulated animal behavior, *i*.*e*. a mushroom body output neuron.

Our data show that the dendritic tree of the MBON is electrotonically compact, despite the complex architecture that includes a high degree of branching. This data is in agreement with a prior electrophysiological characterization of an MBON in locust (Cassenaer and Laurent, 2007) and a similar feature has been reported for neurons in the stomatogastric ganglion of crayfish. Here the electrotonic compactness supports linear integration of synaptic inputs across extensive arborizations and likely serves to functionally compensate for inter-individual variability (Otopalik et al, 2017, 2019). The location of an individual synaptic input within the dendritic tree has therefore only a minor effect on the amplitude of the neuron’s output, despite large variations of local dendritic potentials. This effect was particularly striking when we restricted the analysis to a population of KCs with identical numbers of synapses that all elicited highly stereotypical responses. The compactification of the neuron is likely related to the architectural structure of its dendritic tree. Together with the relatively small size of many central neurons in *Drosophila*, this indicates that in contrast to large vertebrate neurons, local active amplification or other compensatory mechanisms (Rumsey and Abbott, 2006; Sterratt et al, 2012; Froemke et al, 2005; Gidon and Segev, 2009; Schiller et al, 2000; Polsky et al, 2004) may not be necessary to support input normalization in the dendritic tree. In contrast, for axons it has been recently reported that voltage-gated Na^+^ channels are localized in putative spike initiation zones in a subset of central neurons of *Drosophila* (Ravenscroft et al, 2020). In case *in vivo* physiological data quantitatively describing local active currents become available, they can be incorporated into our model to further increase the agreement between model and biological system.

Encoding of odor information and incorporation of memory traces is not performed by individual KCs but by ensembles of KCs and MBONs. Calcium imaging *in vivo* demonstrated that individual odors evoke activity reliably in approximately 3-9% of KCs (Campbell et al, 2013; Honegger et al, 2011). Our simulations of 1,000 independent trials with random sets of 50 KCs, each representing one distinct odor that activates approximately 5% of the KCs innervating the target MBON, demonstrate that such activation patterns robustly elicit MBON activity in agreement with *in vivo* observations of odor-induced activity that elicited robust increases in action potential frequency in MBONs (Hige et al, 2015a; Warth Pérez Arias et al, 2020). The low variability of depolarizations observed in these simulations indicates that information coding by such activation patterns is highly robust. As a consequence, labeled line representations of odor identity are likely not necessary at the level of KCs since relaying information *via* **any** set of ≈50 KCs is of approximately equal efficiency. Such a model is supported by a recent computational study demonstrating that variability in parameters controlling neuronal excitability of individual KCs negatively affects associative memory performance. The authors provide evidence that compensatory variation mechanisms exist that ensure similar activity levels between all odor encoding KC sets to maintain efficient memory performance (Abdelrahman et al, 2021).

Optical recordings of *in vivo* activity of MBONs revealed selective reductions in MBON activity in response to aversive odor training (Jacob and Waddell, 2020; Pai et al, 2013; Owald and Waddell, 2015; Perisse et al, 2016; Séjourné et al, 2011) or to optogenetic activation of selective DANs (Hige et al, 2015a). More generally, both depression and potentiation of MBON activity have been previously observed in different MBON modules *in vivo* (Owald and Waddell, 2015; Hige et al, 2015a; Perisse et al, 2016; Plaçais et al, 2013; Pai et al, 2013; Séjourné et al, 2011; Jacob and Waddell, 2020). In addition, recent studies have observed changes in KC stimulus representations after conditioning (Delestro et al, 2020; Shyu et al, 2019) that may be due to learning-dependent modulations of synaptic PN input to KCs (Baltruschat et al, 2021). Our computational model allowed us to implement and compare these two mechanisms changing MBON output: One is a change in the strength of the KC>MBON synapses (over a range of ±25%); the other is a change in the number of activated KCs (over the same range). Interestingly, we found almost linear relationships between the number of active KCs and the resulting depolarizations, and the same between the strength of synapses and MBON depolarization. Decreasing or increasing either of these variables by 25% significantly altered the level of MBON activation with only minor differences in the extend to which these modifications contributed to MBON depolarization. The two different mechanisms of altering MBON output are potentially utilized for the establishment of different types of memory with fast and local alterations of synaptic transmission likely essential for short-term memory while structural changes may ensure maintenance of memory over long periods of time. In addition, differential modes of plasticity may be required at potentially more static parts of the MB circuitry like the food-related part that is not entirely based on stochastic connectivity (Zheng et al, 2022).

Our simulation data thus shows that the KC>MBON architecture represents a biophysical module that is well-suited to simultaneously process changes based on either synaptic and/or network modulation. Together with the electrotonic nature of the MBONs the interplay between KCs and MBONs thus ensures reliable information processing and memory storage despite the stochastic connectivity of the memory circuitry. While our study focuses on the detailed activity patterns within a single neuron, the availability of large parts of the fly connectome at the synaptic level, in combination with realistic models for synaptic dynamics, should make it possible to extend this work to circuit models to gain a network understanding of the computational basis of decision making.

## 4 Materials and Methods

### 4.1 Fly Stocks

Flies were reared at 25^°^C and 65% humidity on standard fly food. Following stocks were used in this study: 10XUAS-IVS-mCD8::GFP (Pfeiffer et al, 2010) (BDSC 32186), MB082C-Gal4 (Aso et al, 2014b) (BDSC 68286), 2xUAS-EGFP (BDSC 23867) (all from the Bloomington Drosophila Stock Center, Indiana).

### 4.2 Immunohistochemistry

Male flies expressing GFP in MBON-α3 were fixed for 3.5h at 4^°^C in 4% PFA containing PBST (0.2% Triton-X100). Flies were then washed for at least 3 × 30 min at RT in PBST before dissection. Dissected brains were then labelled with primary antibodies (rabbit anti-GFP (A6455, Life technologies, Carlsbad, USA 1:2000) and mouse anti-Brp nc82, Developmental Studies Hybridoma Bank, Iowa, USA, 1:200) for two nights at 4^°^C. After incubation, brains were washed at least 6 × 30 min in PBST. Secondary antibodies (Alexa488 (goatαrabbit) and Alexa568 (goatαmouse) coupled antibodies, Life technologies, Carlsbad, USA, 1:1000) were applied for two nights at 4^°^C. After a repeated washing period, brains were submerged in Vectashield (Vector Laboratories, Burlingame, USA), mounted onto microscope slides, and stored at 4^°^C.

Images were acquired using a LSM 710 confocal scanning microscope with a 25x Plan-NEOFLUAR, NA 0.8 Korr DIC, oil objective (Carl Zeiss GmbH, Jena, Germany) and a Leica STELLARIS 8 confocal microscope with a 20x HC PL APO NA 0.75 multi-immersion objective (Leica Microsystems, Wetzlar, Germany). Raw images were projected with Fiji (Schindelin et al, 2012) and cropped in Photoshop (Adobe, San José, USA). Uniform adjustments of brightness and contrast were performed.

### 4.3 Electrophysiology

All patch clamp recordings were performed at room temperature similar to prior descriptions (Hige et al, 2015a,b). Flies were flipped 1d after hatching to obtain 2-3 d old flies. Due to the ventral location of the MBON-α3 somata within the brain, we performed the recordings *ex vivo*. For preparation, flies expressing GFP in MBON-α3 were briefly anesthetized on ice before removing the entire brain out of the head capsule. The preparation was performed in oxygenated (95% O_2_ and 5% CO_2_) high glucose (260mM) extracellular saline. Analogous to a prior study, the brains were incubated for 30s in 0.5mg°ml protease (from Streptomyces griseus, CAS# 90036-06-0, Sigma-Aldrich, St. Louis, USA) containing extracellular saline (Wilson, 2004). The brain was then transferred to standard extracellular saline (Wilson, 2004) (in mM: 103 NaCl, 3 KCl, 5 TES, 10 trehalose, 260 glucose, 7 sucrose, 26 NaHCO_3_, 1 NaH_2_PO_4_, 1.5 CaCl_2_, 4 MgCl_2_, pH 7.3, 280 − 290mOsmol/kg adjusted with sucrose). For physiological recordings, the brain was transferred to a glass bottom chamber with continuously perfused (2ml/min) oxygenated standard extracellular saline and held in place by a custom made platinum frame.

We used glass capillaries (GB150 (F-)-10P, Science Products, Hofheim, Germany) and a horizontal puller (Flaming Brown Micropipette Puller P-1000, Sutter instruments, Novato, USA) to obtain pipettes with a resistance of 7 − 10M*Ω*. The puller was equipped with a 2.5mm box filament and patch pipettes were pulled with a program consisting of 2 lines (Line 1/ Line 2: Heat 472/463 with a ramp value of 468, Pull 0/41, Velocity 52/70, Delay 1/10, Pressure 350). For recordings, patch pipettes were filled with internal solution (Mauss et al, 2014) (in mM: 140 K-aspartate, 10 HEPES, 4 Mg-ATP, 0.5 Na-GTP, 1 EGTA, 1 KCl, pH 7.29 adjusted with KOH, 265mOsmol°kg). Whole-cell recordings were made using the EPC10 amplifier (HEKA, Reutlingen, Germany) and the PatchMaster software (HEKA, Reutlingen, Germany). Signals were low-pass filtered at 3kHz, and digitized at 10kHz via a digital-to-analog converter. The liquid junction potential of 13mV was corrected online. Patch pipettes were guided under visual control with an upright microscope (BX51WI; Olympus, Tokio, Japan) equipped with a 60x water immersion objective (LUMPlanFl/IR; Olympus, Tokio, Japan) and a CCD camera (Orca 05G, Hamamatsu, Japan). GFP signal of target cells was visualized through fluorescence excitation (100W fluorescence lamp, Carl Zeiss GmbH, Jena, Germany), and a dual-band emission filter set (512/630 HC Dualband Filter GFP/DsRed, AHF Analysentechnik, Germany). We used GFP expression in soma and neurite to identify MBON-α3 neurons. During recordings, the fluorescence excitation was shut off to minimize phototoxic effects. The complete setup was mounted on an air-damped table while being shielded by a Faraday cage. Series resistance was maintained below 90M*Ω* and compensated for up to 35% through the amplifier’s compensation circuitry. In current-clamp mode, cells were held at a baseline of −60mV to measure the relevant parameters for the model. Signals were recorded with a sample rate of 20kHz or 50kHz and low-pass filtered at 5kHz.

To determine passive electrical membrane properties and qualitative patch characteristics, a series of standard protocols were performed:

#### VCstep without compensation

In voltage-clamp mode, a voltage-step (100ms, 5mV) is applied without compensating the series resistance. The corresponding current trace was then used to determine R_series_ and R_input_ using Ohms law and R_m_ was determined as (R_series_ − R_input_)(Numberger and Draguhn, 1996) in Igor Pro (WaveMetrics, Portland, USA).

#### CCdehyper

In current-clamp mode, a step-protocol was performed and recorded at 50kHz. Each sweep starts with 0pA for 10ms, which is followed by a current injection (−26pA to 32pA, with 2pA increments) and a step duration of 400ms. The resulting changes in membrane potential and induced action potentials were then analyzed and plotted with Matlab (MathWorks, Natick, USA). The maximum depolarization was calculated as a mean of the last 70 sample points of the stimulation sweep in each trace. The resting membrane potential was calculated as the mean of the first 500 sample points before the stimulation in each trace. The absolute action potential was measured as the maximum of each trace. The relative action potential amplitude resulted from subtracting the baseline from the absolute value. The baseline resulted from the mean of a 400 sample points window, starting 700 sample points upstream of the maxima in each trace. The resulting trace of the −10pA injection was used to calculate the membrane time constant *τ*_m_ of each cell.

#### CCcontinuous

In current-clamp mode, no current was injected throughout a 60s recording at 50kHz. The resting membrane potential was determined as the mean of the baseline in the recording with Igor Pro and spontaneous activity was plotted with Matlab.

#### CCinject

In current-clamp mode, a short current pulse (−10pA, 200ms) was injected and recorded at 20kHz. We performed 35 iterations per cell. The resulting data of three example cells was used for fitting the membrane kinetics of the model as described previously (Gouwens and Wilson, 2009). Averaging, normalisation (V_m_-V_m0_/V_m0_) and plotting was performed with Matlab. This data was used to perform the model fitting. The baseline (V_m0_) resulted from the average of the sample points before stimulation.

Cells displaying a resting membrane potential higher than −45mV and/or the series resistance was too high (> 90M*Ω*) were excluded from the analysis. Only cells that were firing action potentials were included in the analysis. A list of the five included cells using mCD-GFP as a marker and of the four cells using cytoplasmic GFP as a marker and the corresponding calculated values are provided in Table 1 and Table 1 Source Data 1.

### 4.4 Computational Model

Morphology data for the *Drosophila* MBON-α3-A was originally characterized using scanning electron microscopy (Takemura et al, 2017) and for the present study obtained from the “mushroombody” dataset hosted on the database neuPrint (https://neuprint-examples.janelia.org/; cell ID 54977). These data describe the structure of its dendritic arborization throughout the MBON α3 compartment (Fig. 2B, and Fig. 2 Supplement 1), with coordinates specified at 8 nm pixel resolution and converted to *μ*m for model implementation. The reconstruction is limited to the portion of the neuron in the mushroom body, not including its soma and axon. The geometry of the axon was determined by investigating a separate MBON, MBON14(a3)R (Fig. 2A), obtained from neuPrint’s “hemibrain:v1.1” dataset (https://neuprint.janelia.org/; cell ID 300972942). The axon and synaptic terminal of this neuron were approximated as five separate sections, distinguished by the major branch points identified in the electron microscopic reconstruction (Fig. 2 Supplement 1). Each section was characterized by its total length and average diameter and divided into segments to maintain the average section length of that region in the original reconstruction. They were then appended to the MBON-α3-A proximal neurite as individual linear segments (Fig. 2 Supplement 1). The dimensions of the soma were characterized by confocal microscopy of MBON-α3-A (Fig. 1A). Synaptic locations of the KC innervation of MBON-α3-A were also obtained from neuPrint’s “mushroombody” dataset. MBON-α3-A is innervated by 948 distinct KCs, forming a total of 12,770 synapses with an average of 13.47 synapses per KC. This data set potentially misses up to 10% of synaptic connections and includes errors associated with electron microscopy reconstructions(Takemura et al, 2017).

Electrophysiological properties of the model neuron (Table 2) were determined by fitting the model response to the recorded data, an approach consistent with prior work modeling *Drosophila* central neurons (Gouwens and Wilson, 2009). Somatic voltage excursions in response to 200ms current injections of −10pA were recorded *ex vivo* (Fig. 1 G) and compared with the simulated membrane potential (Fig. 2D). This current injection was replicated *in silico* and the membrane potential change at the soma was fit to our recorded data by varying the cell’s cytoplasm resistivity, specific membrane capacitance, and passive membrane conductance. The fitting was performed using NEURON’s principle axis optimization algorithm, where parameters were tuned until the computed voltage excursion at the soma matched our electrophysiological recordings. The model optimization searched parameter space to minimize an error function defined as the squared differences between the averaged experimental responses to current pulses (Fig. 1 G) and the computed model responses (Fig. 2D-F). The fitting was then verified by normalizing both the model and experimental voltage excursions. The model was first allowed to equilibrate for 30ms simulated time, after which the 200ms current pulse was initiated. The region of data used for the fitting ranged from 32.025ms to 228.03ms, excluding approximately the first and last 2ms of current injection. The domain of the parameter space was restricted to physiologically plausible values: 0.5*μ*F°cm^2^ to 1.5*μ*F°cm^2^ for specific membrane capacitance, 1E − 7S°cm^2^ to 1E − 4S°cm^2^ for passive membrane conductance, and 30*Ω*cm to 400*Ω*cm for cytoplasm resistivity (Table 2). The final values for each parameter are well within these margins. Resting membrane potential and the leak current reversal potential were set at − 55.64mV based on the initial conditions of the recorded data to which the model was fit.

Cholinergic KC to MBON synapses were modeled as localized conductance changes described by alpha functions (Table 2). KC innervation of MBON-α3-A was determined to be cholinergic (Barnstedt et al, 2016). Currents recorded in *Drosophila* KCs were demonstrated to have a reversal potential of 8.9mV and a rise time to a maximal conductance of 0.44ms (Su and O’Dowd, 2003). The maximal conductance for the synapses was set at 1.5627 * 10^−5^*μ*S, the value determined to achieve the target MBON depolarization from monosynaptic KC innervation (Hige et al, 2015b).

All simulations were performed using the NEURON 7.7.2 simulation environment (Hines and Carnevale, 1997) running through a Python 3.8.5 interface in Ubuntu 20.04.1. The morphology data divided the MBON-α3-A dendritic tree into 4336 sections. We added 6 additional sections describing the axon, synaptic terminals, and soma to a total of 4342 sections with an average of 1.24*μ*m in length and 0.29*μ*m in diameter (Table 4). Passive leak channels were inserted along the membrane of the dendrite sections as well as throughout the axon and soma. Each dendritic section was automatically assigned an index number by NEURON and modeled as a piece of cylindrical cable, with connectivity specified by the neuron’s morphology. The coordinate location of each KC synapse was mapped to the nearest section of the MBON dendrite, and the conductance-based synapse model was implemented in the center of that section. Simulations were performed in which each synapse-containing section was individually activated, followed by additional simulations activating groups of synapses corresponding to single or multiple active KCs. A list of all performed simulations related to Figure 5 is provided in Table 3. Membrane potential data was analyzed and plotted with MATLAB R2020b (MathWorks, Natick, USA) and Prism 9 (GraphPad, San Diego, USA).

**Table 4.** Morphological parameters of the model

| Parameter | Value |
| --- | --- |
| number of sections | 4342 |
| average section length | 1.24$\mu\text{m}$ |
| total length | 5377.39$\mu\text{m}$ |
| average diameter | 0.29$\mu\text{m}$ |
| average segment surface area | 1.26$\mu\text{m}^2$ |
| total surface area | 6168.32$\mu\text{m}^2$ |
| soma diameter | 6.45$\mu\text{m}$ |

All code and data files necessary to replicate the simulations are available at: https://doi.org/10.7281/T1/HRK27V.

### 4.5 Statistical Analysis

Statistical analysis and visualization of data was performed in Prism 9 (Graph-Pad, San Diego, USA) and Matlab (MathWorks, Natick, USA). Before a statistical comparison was performed, individual groups were tested in Prism for normality and lognormality with an Anderson-Darling test. Statistical significance was tested with an unpaired parametric t-test or in case of multiple comparisons, with an ordinary corrected parametric one-way ANOVA test. Further information is provided in figure legends.

## Acknowledgments

We would like to thank Glenn Turner and Toshihide Hige for insightful comments on the patch clamp analysis, Hans-Peter Schneider for technical help and Oliver Barnstedt for comments on the manuscript.

---

**Figure 1 Supplement 1.**
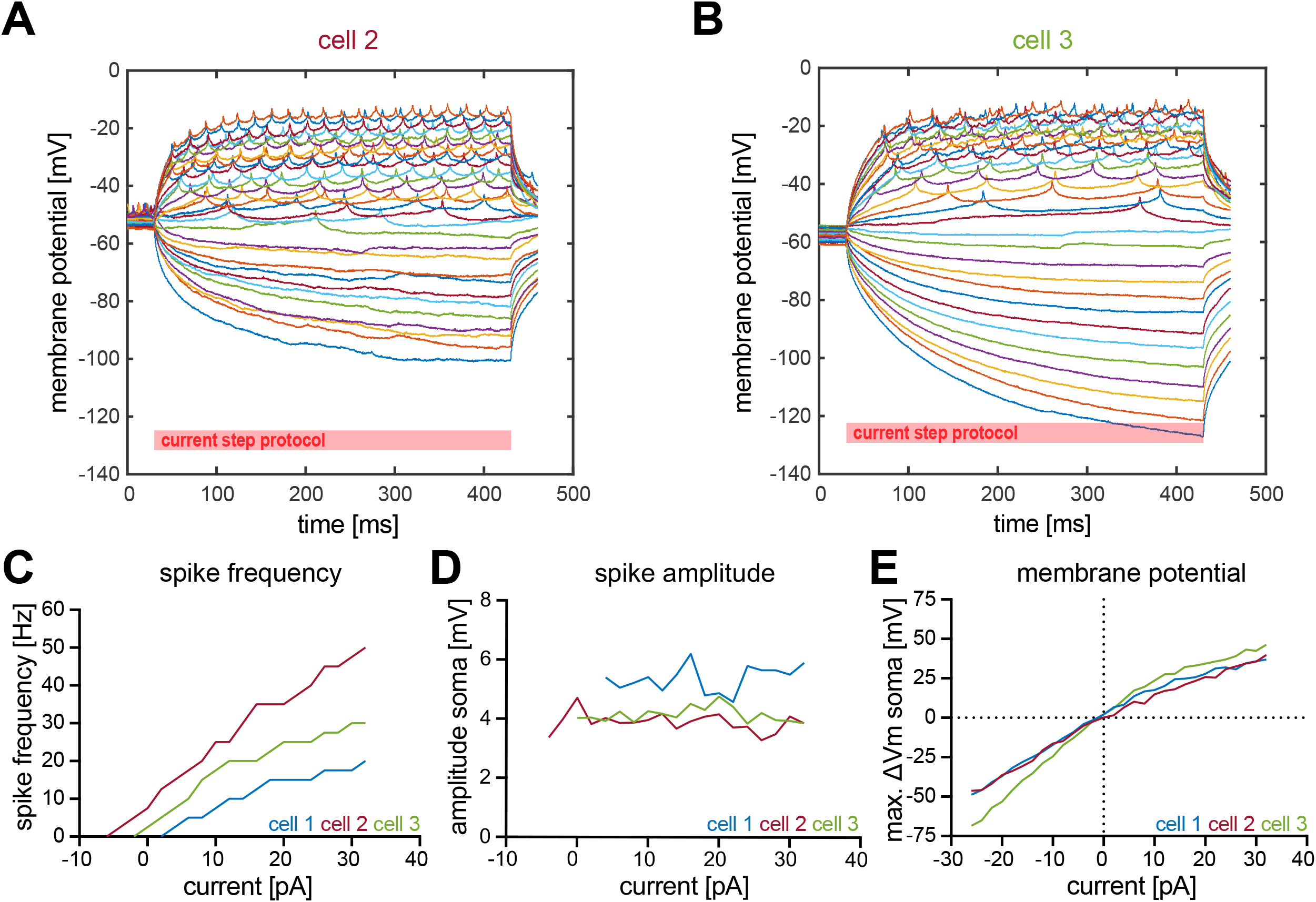
Additional information about the electrophysiology of MBON-α3. Second (A) and third (B) example cells for the evoked neuronal activity of MBON-α3 through a step-wise increasing injection of 400ms current pulses. Pulses start at −26pA (bottom) with increasing 2pA steps and end at +32pA (top). (C) Correlation between frequency of action potential firing and injected current. The frequency describes action potentials per second. (D) Relative change of spike amplitude after current injections. Spike amplitude corresponds to the peak of the highest action potential minus baseline. (E) Change of membrane potential relative to the resting membrane potential after current injection. ΔVm is equal to the maximum depolarisation minus the resting membrane potential.

**Figure 1 Supplement 2.**
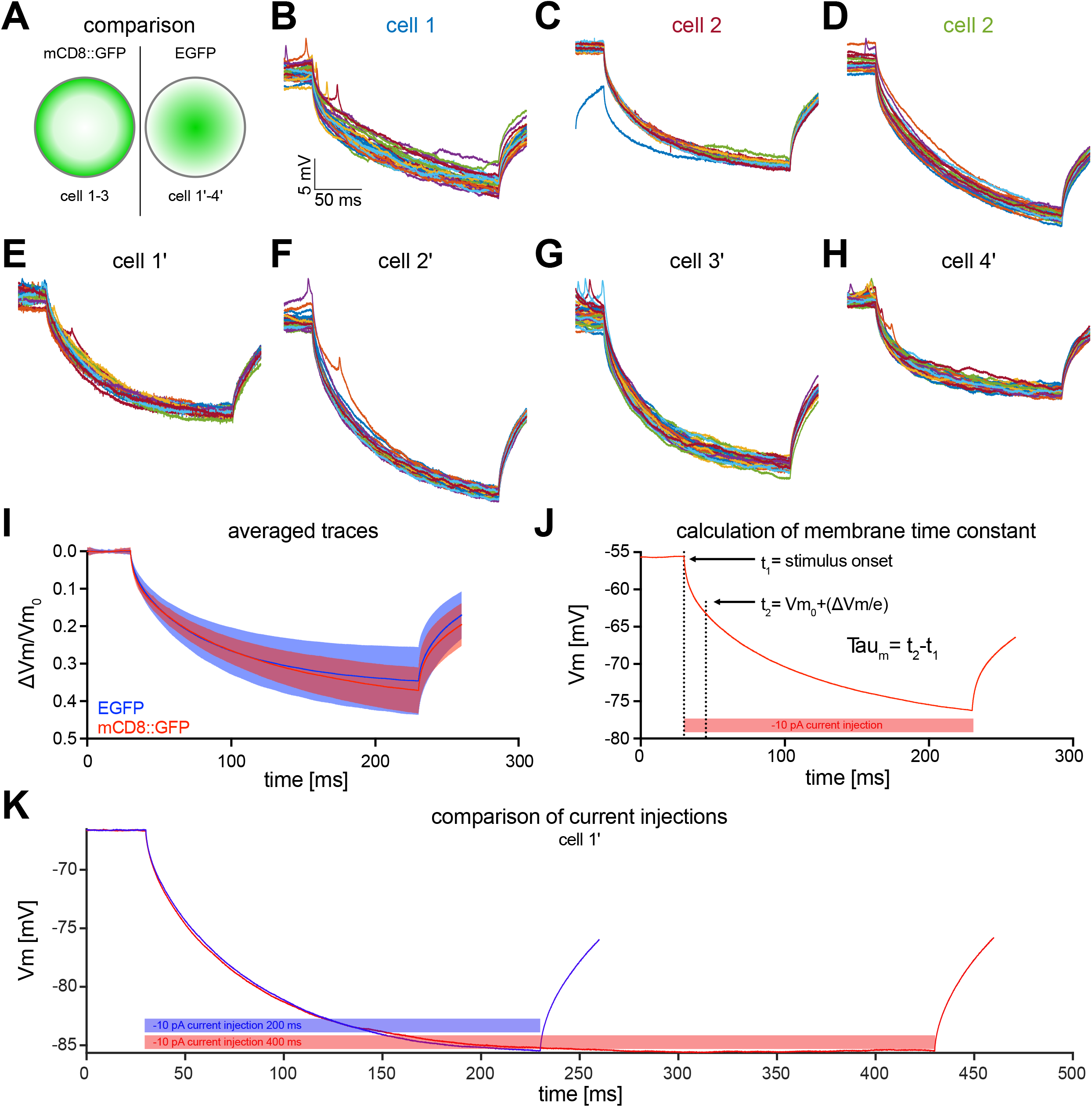
Electrophysiological properties of MBON-α3. (A) MBON-α3 was identified for patch-clamp recordings using either mCD8::GFP (data in B-D) or cytoplasmic EGFP (data in E-H). (B-H) Raw traces of the induced membrane polarization resulting from a 200ms long current injection of 10pA. (I) Comparison of the mean traces of the three cells recorded using mCD8::GFP (B-D, blue) and the four cells recorded using EGFP (E-H, red). Blue and red shadings indicate the SD. (J) Exemplary illustration of the procedure to determine the membrane time constant tau. All parameters used to calculate *τ*_m_ are highlighted. The first vertical dashed line is defined as the time (t_1_) when the current injection starts. The second line (t_2_) is when the voltage has decayed to (Vm(t_1_) + (ΔVm°e)), where ΔVm is the difference between V(t_1_) and the asymptotic voltage, and e is the base of the natural logarithm. Thus, at t_2_, the deflection is at 1/e of the maximum deflection. (K) Comparison of 200ms and 400ms current injections of −10pA from an example cell (cell 1’). No significant difference for *τ*_m_ could be detected between the two stimulation protocols.

**Figure 2 Supplement 1.**
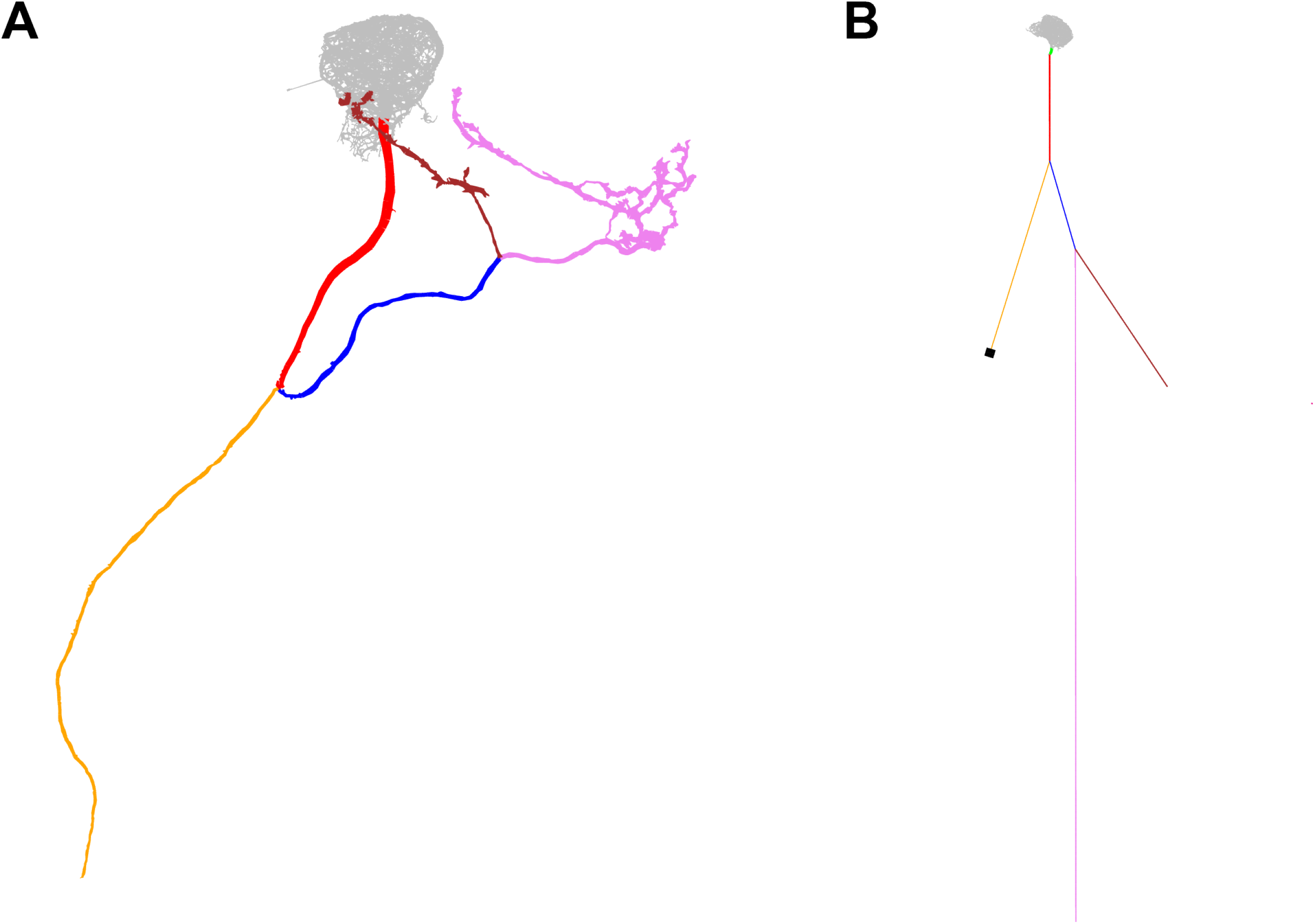
Overview of the axonal sections and soma in the *in silico* model. The pink and brown sections represent the presynaptic release sites of MBON-α3. Color-coded electron-microscopy based reconstruction (A) and *in silico* model (B). The reconstruction of the hemibrain did not include the complete axonal area (yellow) and the soma. (B) illustrates the appended sections and the corresponding diameter and lengths. The soma diameter (black) and axon length (yellow) were estimated from confocal microscopy images and literature.

**Figure 2 Supplement 2.**
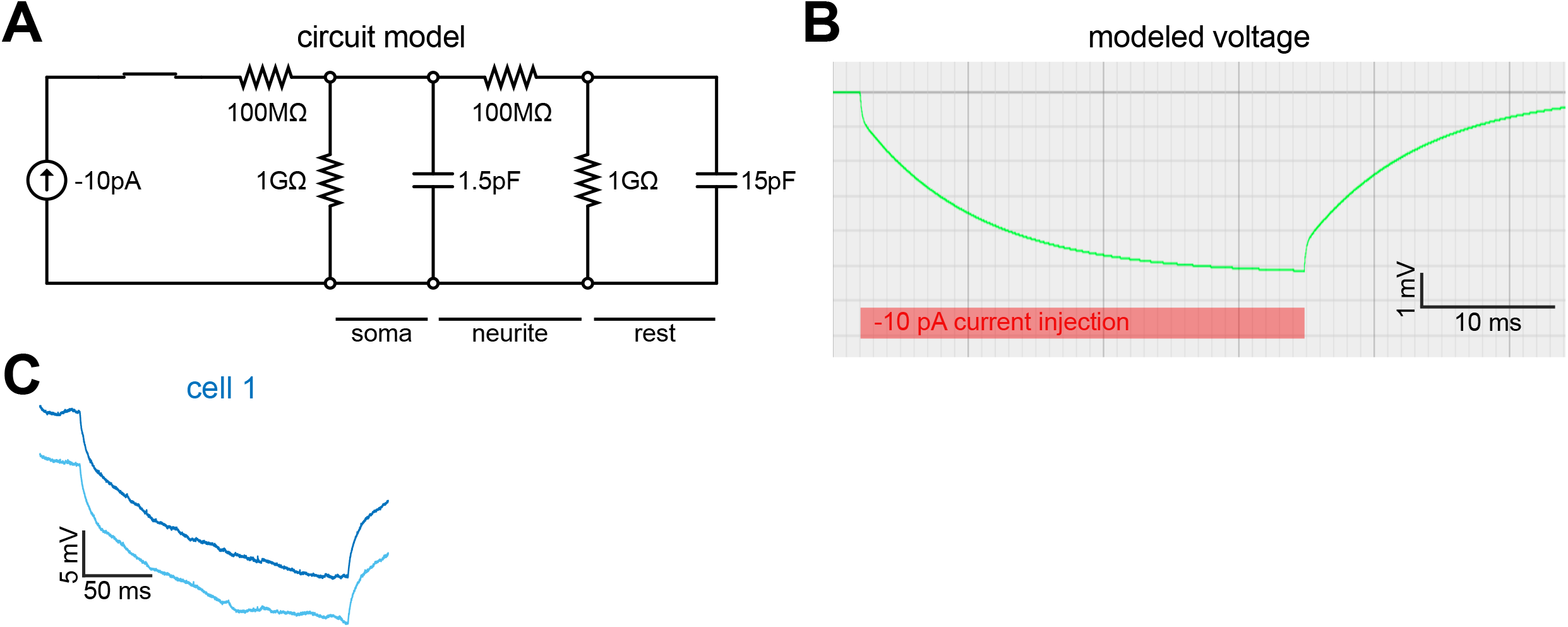
Origin of the fast voltage change followed by slower change subsequent to current steps, visible in Figure 2D,E,F. We hypothesize that this behavior is a reflection of the cell’s morphology, shown in panel A of that Figure, as well as in Figure 2 Supplement 1. In the experiment, as well as in the simulation, current is injected into the soma and recorded there. The soma is a relatively small compartment, with small membrane area. It is connected *via* a long, thin neurite with the rest of the cell which has a substantially larger membrane area. Current injection charges the somatic capacitance very quickly, leading to a rapid change in the somatic voltage at each current step, while the rest of the cell reacts on a much larger time scale. This is visible by the small “kink” right after each change in the injected constant current, Figure 2D. (A) Highly simplified schematic circuit of the cell. The soma is represented by the RC circuit consisting of a 1G*Ω* resistor and a 1.5pF capacitor. The time constant of this circuit is very short, 1.5ms. The neurite connecting it to the rest of the cell (axon and dendrites) is represented by a 100M*Ω* resistor. The rest of the cell is represented by the second RC circuit (1G*Ω* resistor and 15pF capacitor) with a time constant exceeding that of the soma by a factor of ten. (B) Simulated current step protocol in the simplified circuit, with negative current injected for ≈5ms and then removed (red bar). The somatic voltage (green trace) shows the fast change at start and end of the current injection, followed by a slower near-exponential trace. This Link to the circuit simulator allows the reader to simulate the circuit shown in (A). Closing the switch, by clicking on its symbol, starts the −10pA current injection and shows the soma voltage, *i*.*e*. the voltage over the 1.5pF capacitor. Opening the switch (by clicking on the symbol again) results in the circuit returning to its equilibrium state. The voltage trace (green) displays the two characteristic times of the coupled circuit. (C) The two time scales are visible in the electrophysiologically recorded raw (not averaged) voltage, in fact more clearly than in the averaged data in Figure 2D,E,F. Shown here are two representative voltage traces recorded from cell 1 in Figure 1 Supplement 2B.

**Figure 3 Supplement 1.**
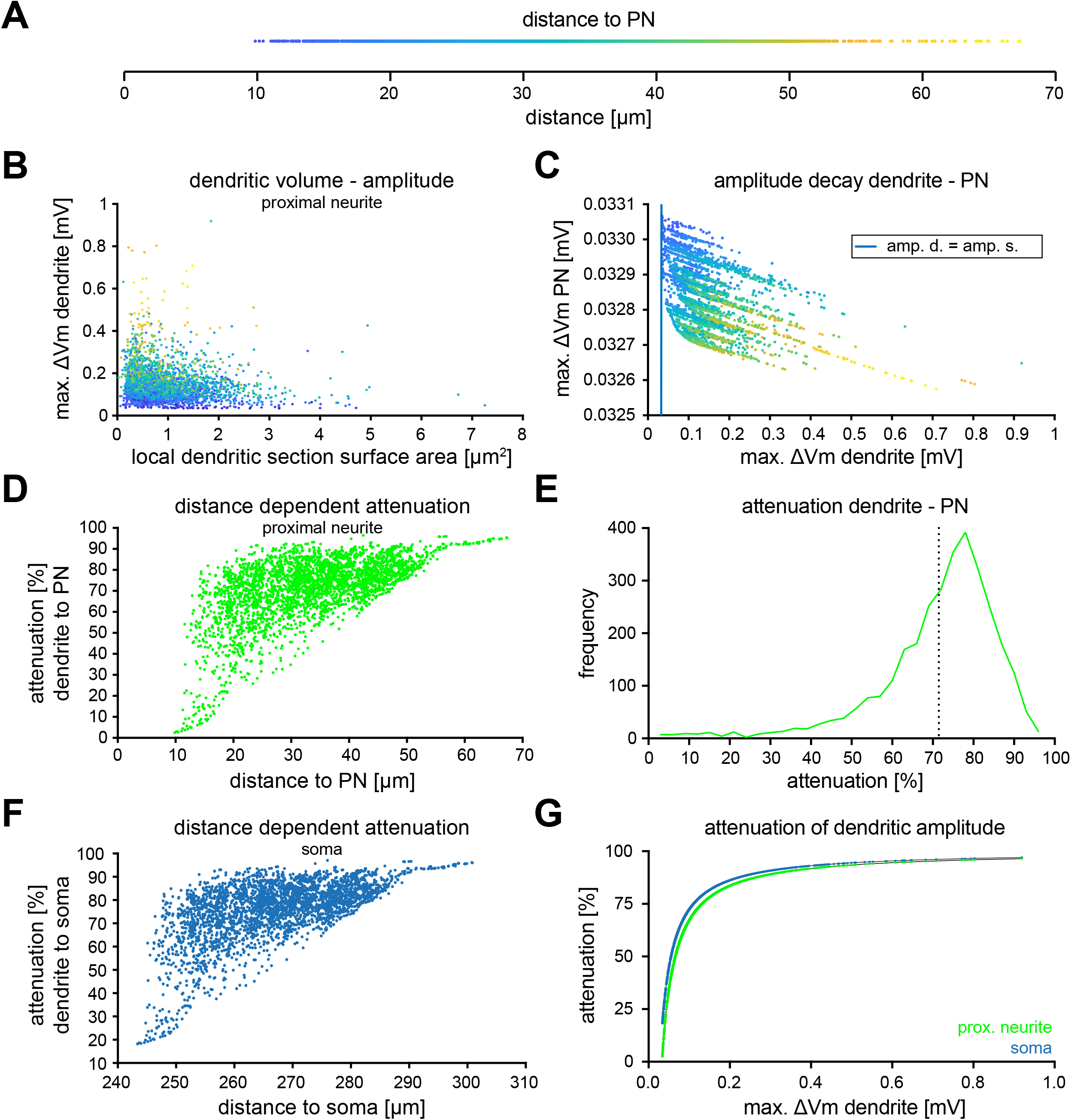
Additional information about voltage attenuation at the proximal neurite and soma. (A) Distance to the proximal neurite for each stimulated dendritic segment with the color code for the following graphs. (B) Scatter plot of the elicited amplitude relative to the resting membrane potential in each stimulated dendritic section as function of its surface area. (C) Scatter plot of the voltage in the proximal neurite as a function of the voltage in the dendrite, both relative to their baseline values. The blue line represents the line of identity. (D) Correlation of the measured distance to the proximal neurite with the percent of voltage attenuation in the proximal neurite for each synaptic section. (E) Distribution of the percentage of voltage attenuation in the soma per synapse. Vertical dotted line represents the mean attenuation in percent. (F) Correlation of the measured distance to the soma with the percent of voltage attenuation in the soma for each synaptic section. (G) Correlation of the attenuation in percent at the soma (blue) or at the proximal neurite (green) relative to the dendritic voltage.

## References

Abdelrahman NY, Vasilaki E, Lin AC (2021) Compensatory variability in network parameters enhances memory performance in the Drosophila mushroom body. Proceedings of the National Academy of Sciences of the United States of America 118(49):e2102158.118. https://doi.org/10.1073/pnas.2102158118

Aso Y, Rubin GM (2016) Dopaminergic neurons write and update memories with cell-type-specific rules. eLife 5:e16.135. https://doi.org/10.7554/eLife.16135

Aso Y, Hattori D, Yu Y, et al (2014a) The neuronal architecture of the mushroom body provides a logic for associative learning. eLife 3:e04.577. https://doi.org/10.7554/eLife.04577

Aso Y, Sitaraman D, Ichinose T, et al (2014b) Mushroom body output neurons encode valence and guide memory-based action selection in Drosophila. eLife 3:e04.580. https://doi.org/10.7554/eLife.04580

Baltruschat L, Prisco L, Ranft P, et al (2021) Circuit reorganization in the Drosophila mushroom body calyx accompanies memory consolidation. Cell Reports 34(11):108,871. https://doi.org/10.1016/j.celrep.2021.108871

Barnstedt O, Owald D, Felsenberg J, et al (2016) Memory-Relevant Mushroom Body Output Synapses Are Cholinergic. Neuron 89(6):1237–1247. https://doi.org/10.1016/j.neuron.2016.02.015

Berger SD, Crook SM (2015) Modeling the Influence of Ion Channels on Neuron Dynamics in Drosophila. Frontiers in Computational Neuroscience 9:139. https://doi.org/10.3389/fncom.2015.00139

Berry JA, Phan A, Davis RL (2018) Dopamine Neurons Mediate Learning and Forgetting through Bidirectional Modulation of a Memory Trace. Cell Reports 25(3):651–662.e5. https://doi.org/10.1016/j.celrep.2018.09.051

Betkiewicz R, Lindner B, Nawrot MP (2020) Circuit and Cellular Mechanisms Facilitate the Transformation from Dense to Sparse Coding in the Insect Olfactory System. eneuro 7(2):ENEURO.0305–18.2020. https://doi.org/10.1523/ENEURO.0305-18.2020

Borst A, Haag J (1996) The intrinsic electrophysiological characteristics of fly lobula plate tangential cells: I. Passive membrane properties. Journal of Computational Neuroscience 3(4):313–336. https://doi.org/10.1007/BF00161091

Bouzaiane E, Trannoy S, Scheunemann L, et al (2015) Two Independent Mushroom Body Output Circuits Retrieve the Six Discrete Components of Drosophila Aversive Memory. Cell Reports 11(8):1280–1292. https://doi.org/10.1016/j.celrep.2015.04.044

Burke CJ, Huetteroth W, Owald D, et al (2012) Layered reward signalling through octopamine and dopamine in Drosophila. Nature 492(7429):433–437. https://doi.org/10.1038/nature11614

Campbell RAA, Honegger KS, Qin H, et al (2013) Imaging a population code for odor identity in the Drosophila mushroom body. The Journal of Neuroscience: The Official Journal of the Society for Neuroscience 33(25):10,568–10,581. https://doi.org/10.1523/JNEUROSCI.0682-12.2013

Caron SJC, Ruta V, Abbott LF, et al (2013) Random convergence of olfactory inputs in the Drosophila mushroom body. Nature 497(7447):113–117. https://doi.org/10.1038/nature12063

Cassenaer S, Laurent G (2007) Hebbian STDP in mushroom bodies facilitates the synchronous flow of olfactory information in locusts. Nature 448(7154):709– 713. https://doi.org/10.1038/nature05973

Cognigni P, Felsenberg J, Waddell S (2018) Do the right thing: Neural network mechanisms of memory formation, expression and update in Drosophila. Current Opinion in Neurobiology 49:51–58. https://doi.org/10.1016/j.conb.2017.12.002

Cohn R, Morantte I, Ruta V (2015) Coordinated and Compartmentalized Neuromodulation Shapes Sensory Processing in Drosophila. Cell 163(7):1742–1755. https://doi.org/10.1016/j.cell.2015.11.019

de Belle J, Heisenberg M (1994) Associative odor learning in Drosophila abolished by chemical ablation of mushroom bodies. Science 263(5147):692–695. https://doi.org/10.1126/science.8303280

Delestro F, Scheunemann L, Pedrazzani M, et al (2020) In vivo large-scale analysis of Drosophila neuronal calcium traces by automated tracking of single somata. Scientific Reports 10(1):7153. https://doi.org/10.1038/s41598-020-64060-x

Eichler K, Li F, Litwin-Kumar A, et al (2017) The complete connectome of a learning and memory centre in an insect brain. Nature 548(7666):175–182. https://doi.org/10.1038/nature23455

Felsenberg J, Jacob PF, Walker T, et al (2018) Integration of Parallel Opposing Memories Underlies Memory Extinction. Cell 175(3):709–722.e15. https://doi.org/10.1016/j.cell.2018.08.021

Froemke RC, Poo Mm, Dan Y (2005) Spike-timing-dependent synaptic plasticity depends on dendritic location. Nature 434(7030):221–225. https://doi.org/10.1038/nature03366

Gidon A, Segev I (2009) Spike-timing-dependent synaptic plasticity and synaptic democracy in dendrites. Journal of Neurophysiology 101(6):3226–3234. https://doi.org/10.1152/jn.91349.2008

Gouwens NW, Wilson RI (2009) Signal Propagation in Drosophila Central Neurons. Journal of Neuroscience 29(19):6239–6249. https://doi.org/10.1523/JNEUROSCI.0764-09.2009

Groschner LN, Malis JG, Zuidinga B, et al (2022) A biophysical account of multiplication by a single neuron. Nature 603(7899):119–123. https://doi.org/10.1038/s41586-022-04428-3

Gruntman E, Turner GC (2013) Integration of the olfactory code across dendritic claws of single mushroom body neurons. Nature Neuroscience 16(12):1821– 1829. https://doi.org/10.1038/nn.3547

Gu H, O’Dowd DK (2006) Cholinergic synaptic transmission in adult Drosophila Kenyon cells in situ. The Journal of Neuroscience: The Official Journal of the Society for Neuroscience 26(1):265–272. https://doi.org/10.1523/JNEUROSCI.4109-05.2006

Handler A, Graham TG, Cohn R, et al (2019) Distinct Dopamine Receptor Pathways Underlie the Temporal Sensitivity of Associative Learning. Cell 178(1):60– 75.e19. https://doi.org/10.1016/j.cell.2019.05.040

Hige T, Aso Y, Modi MN, et al (2015a) Heterosynaptic Plasticity Underlies Aversive Olfactory Learning in Drosophila. Neuron 88(5):985–998. https://doi.org/10.1016/j.neuron.2015.11.003

Hige T, Aso Y, Rubin GM, et al (2015b) Plasticity-driven individualization of olfactory coding in mushroom body output neurons. Nature 526(7572):258–262. https://doi.org/10.1038/nature15396

Hines ML, Carnevale NT (1997) The NEURON Simulation Environment. Neural Computation 9(6):1179–1209. https://doi.org/10.1162/neco.1997.9.6.1179

Honegger KS, Campbell RAA, Turner GC (2011) Cellular-Resolution Population Imaging Reveals Robust Sparse Coding in the Drosophila Mushroom Body. Journal of Neuroscience 31(33):11,772–11,785. https://doi.org/10.1523/JNEUROSCI.1099-11.2011

Jacob PF, Waddell S (2020) Spaced Training Forms Complementary Long-Term Memories of Opposite Valence in Drosophila. Neuron 106(6):977–991.e4. https://doi.org/10.1016/j.neuron.2020.03.013

Lai SL, Lee T (2006) Genetic mosaic with dual binary transcriptional systems in Drosophila. Nature Neuroscience 9(5):703–709. https://doi.org/10.1038/nn1681

Li F, Lindsey JW, Marin EC, et al (2020) The connectome of the adult Drosophila mushroom body provides insights into function. eLife 9:e62.576. https://doi.org/10.7554/eLife.62576

Lin HH, Lai JSY, Chin AL, et al (2007) A Map of Olfactory Representation in the Drosophila Mushroom Body. Cell 128(6):1205–1217. https://doi.org/10.1016/j.cell.2007.03.006

Litwin-Kumar A, Harris KD, Axel R, et al (2017) Optimal Degrees of Synaptic Connectivity. Neuron 93(5):1153–1164.e7. https://doi.org/10.1016/j.neuron.2017.01.030

Livingstone MS, Sziber PP, Quinn WG (1984) Loss of calcium/calmodulin responsiveness in adenylate cyclase of rutabaga, a Drosophila learning mutant. Cell 37(1):205–215. https://doi.org/10.1016/0092-8674(84)90316-7

Mauss AS, Meier M, Serbe E, et al (2014) Optogenetic and Pharmacologic Dissection of Feedforward Inhibition in Drosophila Motion Vision. Journal of Neuroscience 34(6):2254–2263. https://doi.org/10.1523/JNEUROSCI.3938-13.2014

Miyashita T, Kikuchi E, Horiuchi J, et al (2018) Long-Term Memory Engram Cells Are Established by c-Fos/CREB Transcriptional Cycling. Cell Reports 25(10):2716–2728.e3. https://doi.org/10.1016/j.celrep.2018.11.022

Niebur E (2008) Neuronal cable theory. Scholarpedia 3(5):2674. https://doi.org/10.4249/scholarpedia.2674

Numberger M, Draguhn A (1996) Patch-Clamp-Technik. Labor im Fokus, Spektrum, Akad. Verl, Heidelberg

Otopalik AG, Sutton AC, Banghart M, et al (2017) When complex neuronal structures may not matter. eLife 6:e23.508. https://doi.org/10.7554/eLife.23508

Otopalik AG, Lane B, Schulz DJ, et al (2019) Innexin expression in electrically coupled motor circuits. Neuroscience Letters 695:19–24. https://doi.org/10.1016/j.neulet.2017.07.016

Owald D, Waddell S (2015) Olfactory learning skews mushroom body output pathways to steer behavioral choice in Drosophila. Current Opinion in Neurobiology 35:178–184. https://doi.org/10.1016/j.conb.2015.10.002

Owald D, Felsenberg J, Talbot CB, et al (2015a) Activity of Defined Mushroom Body Output Neurons Underlies Learned Olfactory Behavior in Drosophila. Neuron 86(2):417–427. https://doi.org/10.1016/j.neuron.2015.03.025

Owald D, Lin S, Waddell S (2015b) Light, heat, action: Neural control of fruit fly behaviour. Philosophical Transactions of the Royal Society B: Biological Sciences 370(1677):20140,211. https://doi.org/10.1098/rstb.2014.0211

Pai TP, Chen CC, Lin HH, et al (2013) Drosophila ORB protein in two mushroom body output neurons is necessary for long-term memory formation. Proceedings of the National Academy of Sciences 110(19):7898–7903. https://doi.org/10.1073/pnas.1216336110

Perisse E, Owald D, Barnstedt O, et al (2016) Aversive Learning and Appetitive Motivation Toggle Feed-Forward Inhibition in the Drosophila Mushroom Body. Neuron 90(5):1086–1099. https://doi.org/10.1016/j.neuron.2016.04.034

Pfeiffer BD, Ngo TTB, Hibbard KL, et al (2010) Refinement of Tools for Targeted Gene Expression in Drosophila. Genetics 186(2):735–755. https://doi.org/10.1534/genetics.110.119917

Plaçais PY, Trannoy S, Friedrich AB, et al (2013) Two Pairs of Mushroom Body Efferent Neurons Are Required for Appetitive Long-Term Memory Retrieval in Drosophila. Cell Reports 5(3):769–780. https://doi.org/10.1016/j.celrep.2013.09.032

Polsky A, Mel BW, Schiller J (2004) Computational subunits in thin dendrites of pyramidal cells. Nature Neuroscience 7(6):621–627. https://doi.org/10.1038/nn1253

Ravenscroft TA, Janssens J, Lee PT, et al (2020) Drosophila Voltage-Gated Sodium Channels Are Only Expressed in Active Neurons and Are Localized to Distal Axonal Initial Segment-like Domains. The Journal of Neuroscience: The Official Journal of the Society for Neuroscience 40(42):7999–8024. https://doi.org/10.1523/JNEUROSCI.0142-20.2020

Rumsey CC, Abbott LF (2006) Synaptic democracy in active dendrites. Journal of Neurophysiology 96(5):2307–2318. https://doi.org/10.1152/jn.00149.2006

Sakmann B, Neher E (1995) Single-Channel recording. In: Single-Channel Recording. Springer US, Boston, MA, https://doi.org/10.1007/978-1-4419-1229-9

Scheffer LK, Xu CS, Januszewski M, et al (2020) A connectome and analysis of the adult Drosophila central brain. eLife 9:e57.443. https://doi.org/10.7554/eLife.57443

Schiller J, Major G, Koester HJ, et al (2000) NMDA spikes in basal dendrites of cortical pyramidal neurons. Nature 404(6775):285–289. https://doi.org/10.1038/35005094

Schindelin J, Arganda-Carreras I, Frise E, et al (2012) Fiji: An open-source platform for biological-image analysis. Nature Methods 9(7):676–682. https://doi.org/10.1038/nmeth.2019

Séjourné J, Plaçais PY, Aso Y, et al (2011) Mushroom body efferent neurons responsible for aversive olfactory memory retrieval in Drosophila. Nature Neuroscience 14(7):903–910. https://doi.org/10.1038/nn.2846

Shyu WH, Lee WP, Chiang MH, et al (2019) Electrical synapses between mushroom body neurons are critical for consolidated memory retrieval in Drosophila. PLOS Genetics 15(5):e1008.153. https://doi.org/10.1371/journal.pgen.1008153

Siegenthaler D, Escribano B, Bräuler V, et al (2019) Selective suppression and recall of long-term memories in Drosophila. PLoS biology 17(8):e3000.400. https://doi.org/10.1371/journal.pbio.3000400

Sterratt DC, Groen MR, Meredith RM, et al (2012) Spine Calcium Transients Induced by Synaptically-Evoked Action Potentials Can Predict Synapse Location and Establish Synaptic Democracy. PLOS Computational Biology 8(6):e1002.545. https://doi.org/10.1371/journal.pcbi.1002545

Su H, O’Dowd DK (2003) Fast Synaptic Currents in Drosophila Mushroom Body Kenyon Cells Are Mediated by α-Bungarotoxin-Sensitive Nicotinic Acetylcholine Receptors and Picrotoxin-Sensitive GABA Receptors. The Journal of Neuroscience 23(27):9246–9253. https://doi.org/10.1523/JNEUROSCI.23-27-09246.2003

Takemura Sy, Aso Y, Hige T, et al (2017) A connectome of a learning and memory center in the adult Drosophila brain. eLife 6:e26.975. https://doi.org/10.7554/eLife.26975

Tobin WF, Wilson RI, Lee WCA (2017) Wiring variations that enable and constrain neural computation in a sensory microcircuit. eLife 6:e24.838. https://doi.org/10.7554/eLife.24838

Tomchik SM, Davis RL (2009) Dynamics of Learning-Related cAMP Signaling and Stimulus Integration in the Drosophila Olfactory Pathway. Neuron 64(4):510– 521. https://doi.org/10.1016/j.neuron.2009.09.029

Turner GC, Bazhenov M, Laurent G (2008) Olfactory Representations by Drosophila Mushroom Body Neurons. Journal of Neurophysiology 99(2):734– 746. https://doi.org/10.1152/jn.01283.2007

Warth Pérez Arias CC, Frosch P, Fiala A, et al (2020) Stochastic and Arbitrarily Generated Input Patterns to the Mushroom Bodies Can Serve as Conditioned Stimuli in Drosophila. Frontiers in Physiology 11:53. https://doi.org/10.3389/fphys.2020.00053

Wilson RI (2004) Transformation of Olfactory Representations in the Drosophila Antennal Lobe. Science 303(5656):366–370. https://doi.org/10.1126/science.1090782

Yamagata N, Ichinose T, Aso Y, et al (2015) Distinct dopamine neurons mediate reward signals for short- and long-term memories. Proceedings of the National Academy of Sciences 112(2):578–583. https://doi.org/10.1073/pnas.1421930112

Zheng Z, Li F, Fisher C, et al (2022) Structured sampling of olfactory input by the fly mushroom body. Current biology: CB 32(15):3334–3349.e6. https://doi.org/10.1016/j.cub.2022.06.031

